# PRDM3/16 Regulate Chromatin Accessibility Required for NKX2-1 Mediated Alveolar Epithelial Differentiation and Function

**DOI:** 10.1101/2023.12.20.570481

**Authors:** Hua He, Sheila M. Bell, Ashley Kuenzi Davis, Shuyang Zhao, Anusha Sridharan, Cheng-Lun Na, Minzhe Guo, Yan Xu, John Snowball, Daniel T. Swarr, William J. Zacharias, Jeffrey A. Whitsett

**Affiliations:** Key Laboratory of Birth Defects and Related Disease of Women and Children of MOE, West China Second University Hospital Sichuan University, Chengdu, Sichuan, 610041, China; NHC Key Laboratory of Chronobiology, Sichuan University, Sichuan 610041, China; Perinatal Institute, Division of Neonatology and Pulmonary Biology, Cincinnati Children’s Hospital Medical Center; Department of Pediatrics, University of Cincinnati College of Medicine

**Keywords:** PRDM, Alveolar Epithelial Cell, Differentiation, Pulmonary Surfactant, Chromatin Accessibility

## Abstract

Differential chromatin accessibility accompanies and mediates transcriptional control of diverse cell fates and their differentiation during embryogenesis. While the critical role of NKX2-1 and its transcriptional targets in lung morphogenesis and pulmonary epithelial cell differentiation is increasingly known, mechanisms by which chromatin accessibility alters the epigenetic landscape and how NKX2-1 interacts with other co-activators required for alveolar epithelial cell differentiation and function are not well understood. Here, we demonstrate that the paired domain zinc finger transcriptional regulators PRDM3 and PRDM16 regulate chromatin accessibility to mediate cell differentiation decisions during lung morphogenesis. Combined deletion of *Prdm3* and *Prdm16* in early lung endoderm caused perinatal lethality due to respiratory failure from loss of AT2 cell function. *Prdm3/16* deletion led to the accumulation of partially differentiated AT1 cells and loss of AT2 cells. Combination of single cell RNA-seq, bulk ATAC-seq, and CUT&RUN demonstrated that PRDM3 and PRDM16 enhanced chromatin accessibility at NKX2-1 transcriptional targets in peripheral epithelial cells, all three factors binding together at a multitude of cell-type specific cis-active DNA elements. Network analysis demonstrated that PRDM3/16 regulated genes critical for perinatal AT2 cell differentiation, surfactant homeostasis, and innate host defense. Lineage specific deletion of PRDM3/16 in AT2 cells led to lineage infidelity, with PRDM3/16 null cells acquiring partial AT1 fate. Together, these data demonstrate that NKX2-1-dependent regulation of alveolar epithelial cell differentiation is mediated by epigenomic modulation via PRDM3/16.

**Graphical Abstract:** 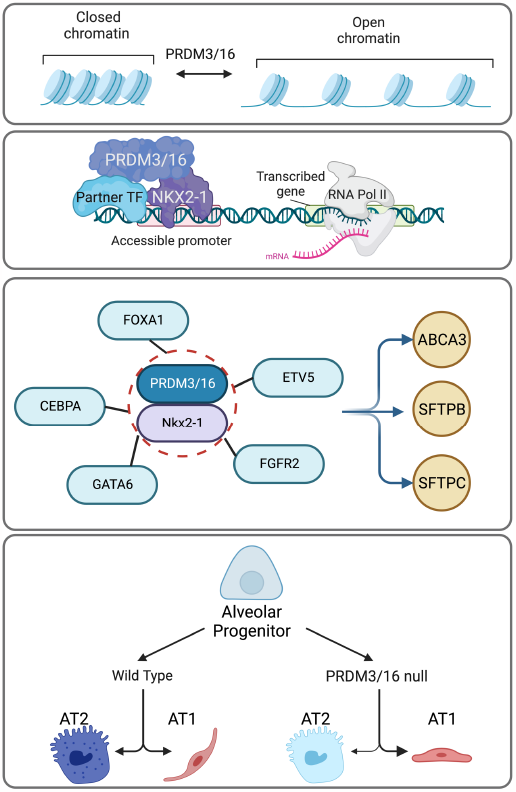

*Model of the role of PRMD3/16 in alveolar development:* PRMD3/16 participate in cell fate specification in the lung by modulating chromatin accessibility (top row) and by partnering with NKX2-1 and partner transcription factors to drive gene expression (second row) via a gene regulatory network required for terminal cell differentiation and surfactant expression in AT2 cells (third row). Loss of PRDM3/16 activity in lung endoderm leads to reduced AT2 quorum, failure of AT2 surfactant function, and transition to an immature AT1 phenotype (bottom panel).

## Introduction

Lung morphogenesis and function are orchestrated by sequential interactions among diverse progenitor cells from the foregut endoderm and splanchnic mesenchyme. Signaling and transcription programs directing cell proliferation and cell fate differentiation are required to place precise numbers of specific cell types in appropriate positions to enable pulmonary function at birth.^1-4^ Cell fate decisions for each of the many pulmonary cell types are instructed by cell-specific gene regulatory networks, which are dynamically used during morphogenesis to control transcription factor activity regulating the expression of a multitude of genes and proteins mediating cell-specific functions. Formation of the mammalian lung foregut endoderm is dependent on transcriptional programs controlled by NKX2-1 (aka thyroid transcription factor 1) and its interactions with other transcription factors to form the conducting airways, peripheral lung saccules, and ultimately, the alveolar regions of the lung required for gas exchange after birth.^5-12^ NKX2-1 is expressed at variable levels in conducting and peripheral lung tubules and is required for branching morphogenesis. SOX2 defines progenitor cells forming the proximal tubules, while epithelial cells in the peripheral acinar buds are marked by expression of SOX9 and ID2.^13-16^ NKX2-1 plays distinct roles in regulating epithelial cell fate during early lung morphogenesis as alveolar type 2 (AT2) and alveolar type 1 (AT1) cells differentiate from SOX9 progenitors. AT1 cell differentiation is directed by the interactions of NKX2-1 with the YAP/TAZ-TEAD family of transcription factors, distinguishing AT1 cell fate choices from AT2 cell fates early in lung development.^10,12,17,18^ NKX2-1 is required for the induction of a diversity of transcriptional targets in AT2 cells to induce pulmonary surfactant lipid, surfactant protein, and innate host defense protein synthesis prior to birth.^1,19^ Incomplete AT2 cell differentiation in preterm infants is associated with surfactant deficiency and causes respiratory distress syndrome (RDS), a major cause of infant morbidity and mortality.^20^ NKX2-1 directly regulates the expression of genes critical for lung function at birth, including those encoding surfactant proteins A, B, C, and D, and genes controlling AT2 cell lipid homeostasis including *Abca3, Scd1*, and *Sreb*. ^8,21,22^ NKX2-1 functions in gene regulatory networks with a number of transcription factors expressed in pulmonary epithelial cells, including FOXA family members (FOXA1 and FOXA2), KLF5, GATA6, TEAD, SREBF1, NFl, and CEBPα, which bind cis-active regulatory elements to activate transcriptional targets critical for surfactant homeostasis prior to and after birth.^7,10,21,23-25^

Accessibility of transcription factors to their targets is regulated by the dynamic control of chromatin structure that is influenced by a diversity of histones and their post-translational modifications.^26^ While the identity and roles of AT2 cell selective transcription factors required for perinatal AT2 cell differentiation and function are increasingly understood, how chromatin is remodeled to provide access to the transcription factors critical for AT2 and AT1 cell functions prior to birth is poorly understood. Remarkably, the dramatic increase in the expression of NKX2-1 transcriptional targets before birth is not accompanied by concomitant increased expression of NKX2-1, supporting the concept that changes in chromatin accessibility and its interactions with a diversity of co-activators influence the activity of NKX2-1 at its transcriptional targets.^8,12,27^

PR(SET) Domain (PRDM) proteins are a family of 19 distinct zinc finger proteins that serve as transcription factors and co-activators known to remodel chromatin and play diverse roles in organogenesis and stem cell functions in multiple tissues. The semi-redundant proteins PRDM3 (also known as EVI1 or MECOM) and PRDM16 (also known as MEL1) share similar structures, expression patterns, and functions.^28^ Both contain a PR domain that has histone methyltransferase activity that may regulate the epigenetic states of chromatin.^29,30^ Multiple zinc finger domains mediate DNA binding.^28^ PRDM3 and PRDM16 play diverse roles in regulating gene expression, with both repressor and activator functions.^29-33^ PRDM16 is required for normal development of hematopoietic stem cells, chondrocytes, adipocytes, maintenance of neural and intestinal stem cells and cardiac morphogenesis.^32,34-38^ PRDM3 and PRDM16 play distinct and opposing roles during craniofacial development.^29,30,35^

The present work evaluated the developmental function of PRDM3 and 16 in lung epithelial cells during lung morphogenesis and identified complementary and cell-type specific roles of PRDM3 and PRDM16 in alveolar epithelial cell differentiation. Deletion of PRDM3 and PRDM16 in the mouse lung endoderm enhanced AT1 cell and inhibited AT2 cell lineage decisions in the embryonic lung and blocked prenatal induction of AT2 cell selective genes critical for surfactant lipid and surfactant-associated protein production required for postnatal survival. Transcriptomic and epigenetic analyses identified gene regulatory networks implicating PRDM3, PRDM16, and NKX2-1 as critical regulators of lung epithelial formation and function, and demonstrate that PRDM3, PRDM16, and NKX2-1 bind at cis-regulatory elements to modulate the expression of AT2 and AT1 selective genes before birth. Therefore, PRDM3/16 function is required to define alveolar epithelial cell fate during embryonic lung morphogenesis and mediate AT2 cell maturation required for the transition to air breathing at birth.

## Results

### Prdm3, Prdm16, and NKX2-1 are co-expressed in lung endoderm

PRDM3 and PRDM16 share similar protein structures and are widely expressed during embryogenesis.^39^ *Prdm3* mRNA and PRDM16 immunofluorescence staining were present in fetal mouse lung epithelial cells as early as E11.5. Thereafter, PRDM3 and 16 were co-localized with NKX2-1 in both peripheral acinar tubules and conducting airways from E16.5 to E18.5 (Figure 1, Figure S1). *Prdm3, Prdm16*, and *Nkx2-1* mRNA levels did not change dynamically during fetal lung development, while the expression of known NKX2-1 transcriptional targets increases markedly prior to birth (Figure S2A).^27,40^

**Figure 1:**
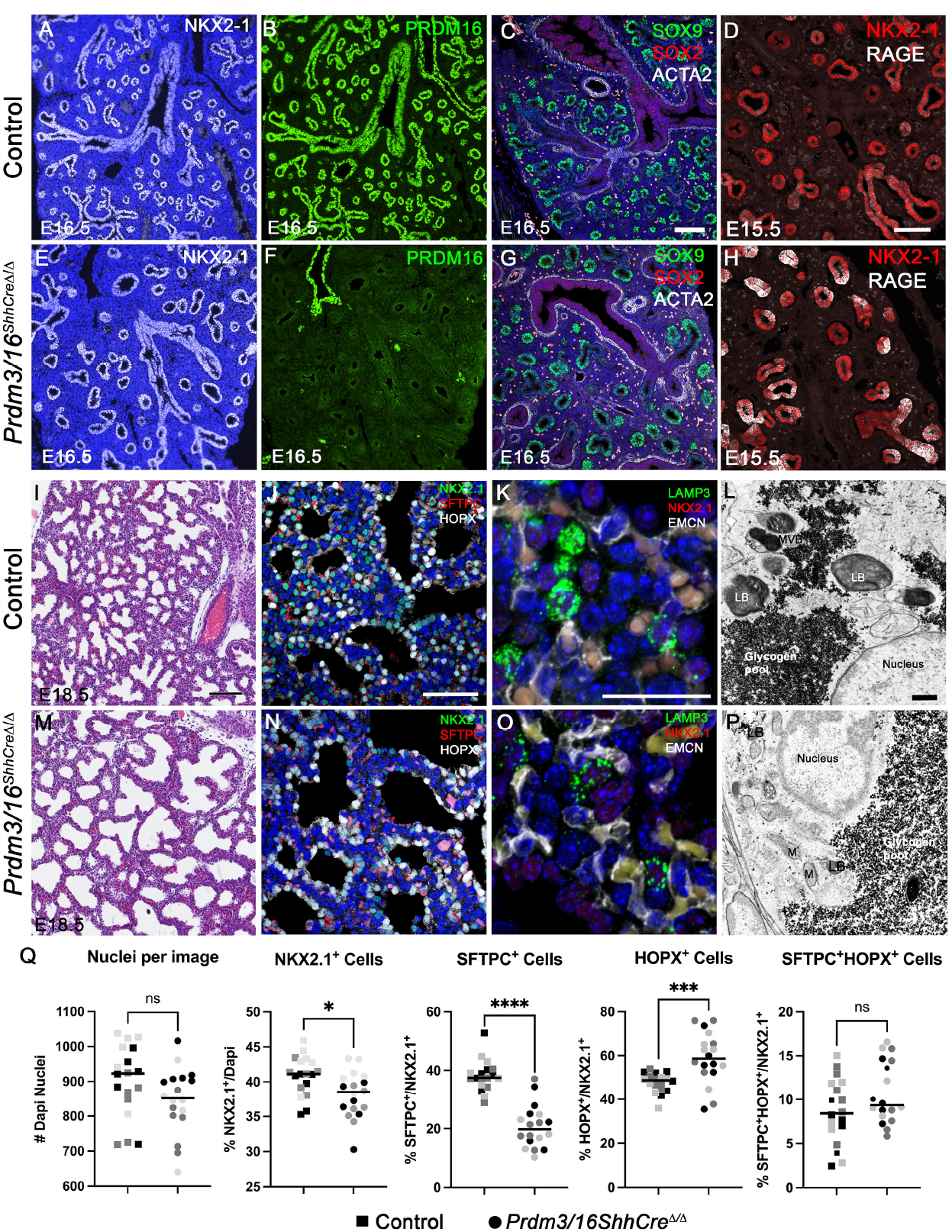
Decreased AT2 cell numbers and differentiation after deletion of Prdm3/16. Immunofluorescence staining of embryonic lung indicates normal expression of NKX2-1after deletion (A, E), loss of PRDM16 staining in lung epithelium in *Prdm3/16*^*ShhCreΔ/Δ*^ fetuses and retention in vascular smooth muscle (B, F), and normal proximal (SOX2+) and distal (SOX9+) epithelial patterning (C, G). At E15.5, the AT1 cell marker RAGE is increased in *Prdm3/16*^*ShhCreΔ/Δ*^ lung (D, H). (I, M) Hematoxylin and eosin staining of E18.5 lung demonstrating poor sacculation in *Prdm3/16*^*ShhCreΔ/Δ*^. (J, K, N, O) Immunofluorescence staining for SFTPC and LAMP3 identifies AT2 cells; AT1 cells are stained for HOPX demonstrating paucity of AT2 cells and reduced LAMP3 expression. (L, P). Electron microscopy of E18.5 lung demonstrates absence of mature lamellar bodies in the *Prdm3/16*^*ShhCreΔ/Δ*^ AT2 cells. (Q) Quantification of AT2, AT1, and AT2/AT1 cell numbers at E18.5 from 3 control and 3 *Prdm3/16*^*ShhCreΔ/Δ*^ fetuses. P-values were calculated using a 2-tailed Mann-Whitney test *p<0.0129, ***p<0.0004, ****p<0.0001, not significant (n.s.). Scale bars in C, D, I (100 mm), J (50 mm), K (25 mm), L (1 mm).

### Prdm3 and Prdm16 are required for cellular differentiation and lung function at birth

To test the roles of PRDM3 and PRDM16 in lung morphogenesis and function, we conditionally deleted each gene individually and in combination using *Shh-Cre* to cause recombination of the *floxed* alleles in lung endoderm, producing *Prdm3*^*ShhCreΔ/Δ*^, *Prdm16*^*ShhCreΔ/Δ*^, and *Prdm3/16*^*ShhCreΔ/Δ*^ mice. *Prdm3*^*ShhCreΔ/Δ*^ *and Prdm16*^*ShhCreΔ/Δ*^ mice survived after birth without visible anatomical defects. RNA sequencing of the sorted lung epithelial cells from *Prdm3/16*^*ShhCreΔ/Δ*^ transgenic mice demonstrated the loss of exon 9 in *Prdm16* RNA and the loss of exon 4 in *Prdm3* RNA (Figure S2B), consistent with deletion of the *floxed* regions in both alleles.^41,42^ Immunofluorescence analysis demonstrated the absence of PRDM16 protein in E16.5 *Prdm3/16*^*ShhCreΔ/Δ*^ lung epithelial cells while staining persisted in the nontargeted endothelial cell population (Figure 1F).

While all newborn *Prdm3/16*^*ShhCreΔ/Δ*^ mice died soon after birth, the only physical difference notable in *Prdm3/16*^*ShhCreΔ/Δ*^ fetuses was the rudimentation of the fifth digit of each paw. While similar in body weight, lung-to-body weight ratios were moderately decreased in both *Prdm3/16*^*ShhCreΔ/Δ*^ and *Prdm3*^*Δ/Δ*^*/16*^*ShhCreΔ/wt*^ fetuses at E18.5 (Figure S2C). At E16.5 and E18.5, while the trachea and esophagus separated normally, peripheral regions of the developing lung were poorly sacculated and septal walls were thickened in *Prdm3/16*^*ShhCreΔ/Δ*^ fetuses (Figure 1). NKX2-1 expression was normally distributed throughout the pulmonary epithelium regardless of PRMD3/16 expression (Figure 1A, E, J, N). In *Prdm3/16*^*ShhCreΔ/Δ*^ E16.5 lungs, SOX2 staining was normally distributed in conducting airways surrounded by αSMA-expressing smooth muscle cells, and SOX9 expression was restricted to peripheral epithelial progenitors, demonstrating preserved cephalo-caudal patterning of the lung (Figure 1C, G). In E18.5 lungs, immunofluorescence demonstrated a paucity of SFTPC^+^ AT2 cells and reduced LAMP3 in *Prdm3/16*^*ShhCreΔ/Δ*^ animals (Figure 1J, K, N, O). HOPX staining was maintained in AT1 cells in *Prdm3/16*^*ShhCreΔ/Δ*^ fetuses. AT2 and AT1 cells were quantified by SFTPC and HOPX staining, demonstrating a decreased proportion of AT2 cells and an increased proportion of AT1 cells in *Prdm3/16*^*ShhCreΔ/Δ*^ (Figure 1Q). Ultrastructural analysis demonstrated a lack of lamellar bodies, the lipid-rich storage sites of pulmonary surfactant in *Prdm3/16*^*ShhCreΔ/Δ*^ AT2 cells (Figure 1L, P). This absence of lamellar bodies and failure of postnatal survival of *Prdm3/16* deficient mice are consistent with respiratory failure caused by the lack of pulmonary surfactant and/or pulmonary hypoplasia.

Recent lineage tracing studies demonstrated that AT1 and AT2 lineage decisions from SOX9 epithelial progenitors occur early in embryonic lung morphogenesis.^43,44^ Therefore, the changes in AT1 and AT2 cell allocation observed in *Prdm3/16*^*ShhCreΔ/Δ*^ mice are likely attributable to roles of PRDM3/16 in lineage decisions occurring early in embryonic lung formation. Consistent with this idea, immunofluorescence staining from AGER was increased in acinar regions of the *Prdm3/16*^*ShhCreΔ/Δ*^ lung tubules as early as E15.5 (Figure 1D, H).

### PRDM3/16 regulates alveolar epithelial cell numbers and differentiation

To test the roles of PRDM3/16 in the regulation of epithelial cell numbers and gene expression, we performed single cell RNA sequencing on *Prdm3/16*^*ShhCreΔ/Δ*^ and control lungs at E18.5. Cell type assignments were made on the basis of interactive clustering and automated cell type annotations using the recently released LungMAP CellRef.^45^ We evaluated 17,581 cells comprising 29 distinct cell clusters expressing cell-type selective RNAs (Figure 3A, Figure S3A). Integrated analysis demonstrated consistent cell type selective gene expression signatures in control and *Prdm3/16*^*ShhCreΔ/Δ*^ samples (Figure S3B), indicating that major cell identity genes were preserved in the absence of PRDM3/16. Consistent with the IHC, scRNA-seq demonstrated a reduction in the proportion of AT2 cells and an increase in AT1 cells after the deletion of *Prdm3* and *16* (Figure 3A, B). While here we have prioritized the analysis of alveolar epithelial cells, these data indicated increased proportions of airway epithelial cell subtypes, including PNECs and secretory cells within the epithelial compartment of *Prdm3/16*^*ShhCreΔ/Δ*^ lungs (Figure 3A, B). Despite the changes in the epithelial compartment, the distribution of cell types within the endothelial and mesenchymal compartments was unchanged (Figure S3C). We compared *Prdm3/16*^*ShhCreΔ/Δ*^ and control AT1 and AT2 cells and identified 198 (AT1) and 207 (AT2) differentially expressed genes (Figure S3D, Table S1). Within the AT2 cell population, expression of genes involved in surfactant homeostasis (*Sftpc, Sftpd*, and *Abca3*), lipid biosynthetic processes (*Scd1, Fabp5, Pi4k2b, Lpcat1, Insig1, Lpin2, Abcd3, and Gpam*), lysosomal function (*Lamp3*), and transmembrane transport (*Atp6v1g, Slc31a1, Slc34a2, Cftr, Atp1b1, and Lcn2*) were all decreased. These data are consistent with decreased ABCA3, LPCAT1, SCD1, and LAMP3 protein detectable by IHC in PRDM3/16-deficient AT2 cells (Figure 1O, 2C, 2E). Within mutant AT1 cells, we noted increased expression of genes associated with translation and regulation of cell migration, whereas downregulated genes were associated with cell adhesion, circulatory system development, and positive regulation of cell motility (Figure S3E). Expression of genes associated with AT1 cell identity and function was generally maintained, including *Ager* and *Hopx* (Figure 2D). Expression of a subset of genes identified as NKX2-1 targets in AT1 cells was repressed in *Prdm3/16*^*ShhCreΔ/Δ*^ AT1 cells including *Cldn18, Aqp5, Pdpn, Samhd1, Pmp22, Gde1, Fbln5, Slco3a1, Gnb4, Bmp4, Akap5, Lmo7*, and *Matn4* (Figure 2D, TableS1) suggesting incomplete terminal differentiation.^11^ Protein expression mirrored these expression changes; immunofluorescence staining of PDPN was decreased, whereas the expression of AQP5 and RAGE was similar in controls and *Prdm3/16*^*ShhCreΔ/Δ*^ at E18.5 (Figure 2E).

**Figure 2.**
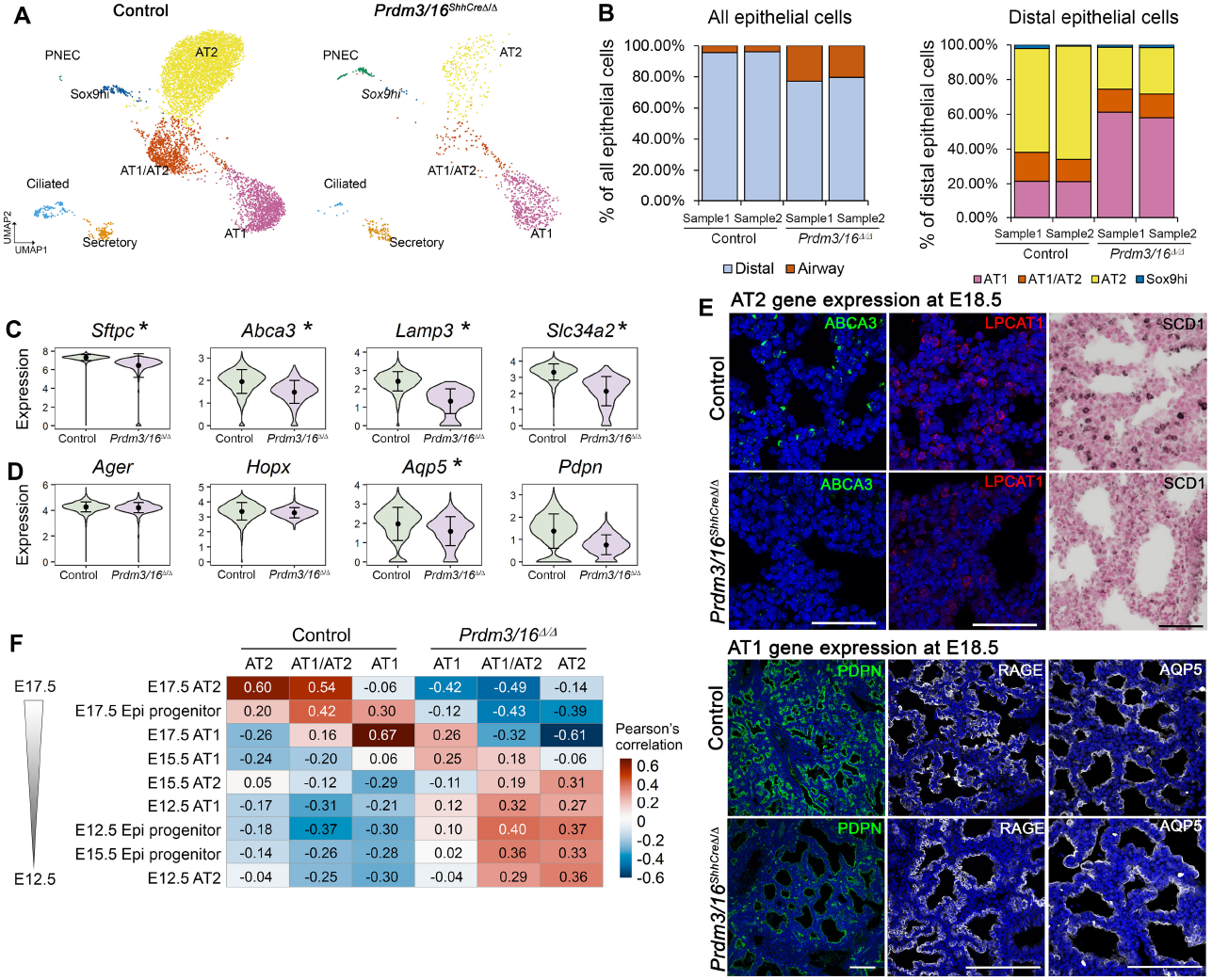
Single-cell RNA-seq (scRNA-seq) analysis of cellular and gene expression alterations in *Prdm3/16*^*ShhCre*D/D^ mouse lung at E18.5. (A) UMAP plots of mouse epithelial cell subsets. (B) Alterations in cell type proportions within all epithelial (left panel) and within distal epithelial (right panel) cells. scRNA-seq data of epithelial cells in (A) were used for the cell type proportion calculations. (C) Violin plot visualization of representative AT2 associated RNAs. (D) Violin plot visualization of representative AT1 associated RNAs. In (C&D), black dots and error bars represent mean±SD; * represents *p*-value of two-tailed Wilcoxon rank sum test ≤0.05, fold change ≥1.5, and expression percentage ≥20%. (E) Immunofluorescence staining of differentially expressed AT1 and AT2 genes in E18.5 lungs (F) Pseudo-bulk correlation analysis with an independent mouse lung developmental time course scRNA-seq data (GSE149563) showing that alveolar epithelial cells from *Prdm3/16*^*ShhCre*Δ/Δ^ mouse lungs are most similar to cells from earlier time points.

**Figure 3.**
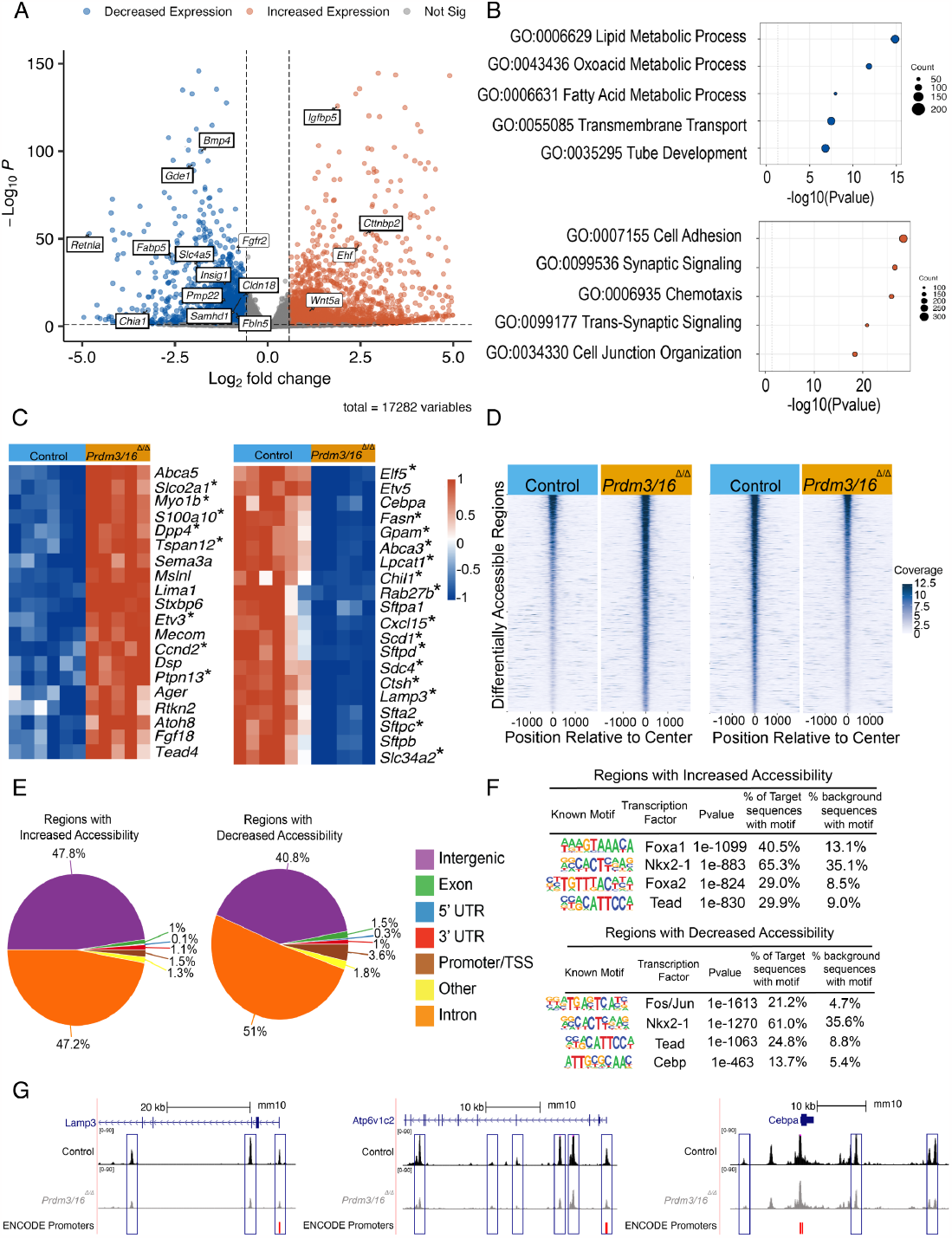
Loss of *Prdm3/16* influences cell differentiation and chromatin accessibility, leading to skewed AT1 and AT2 populations. (A-C) Bulk RNA-Seq analysis of sorted EpCAM^+^ epithelial cells from E17.5 Control and *Prdm3/16*^*ShhCreΔ/Δ*^ (*Prdm3/16*^*Δ/Δ*^) lungs. DESeq2 was used for differential expression analysis utilizing standard cutoffs of log_2_FoldChange > |0.58| and p-value <0.05. (A) Volcano plot showing 1,438 genes with decreased expression and 2,124 genes with increased expression, highlighting genes that are associated with epithelial cell development and mis-regulated in AT1 and AT2 cells, note darkly boxed genes were also observed in the scRNA-seq data. (B) Functional enrichment of gene sets with either increased expression or decreased expression using ToppFun and selecting highly enriched GO: Biological Processes. (C) Heatmaps of normalized gene expression of AT1 and AT2 associated genes, showing an increase in genes associated with AT1 cells and a decrease in genes associated with AT2 cells, a reflection of cell type population size. Asterisk (*) denotes statistical change in both bulk RNAseq and single cell RNA-seq. (D) Heatmaps of ATAC-seq data made with the R package tornado plot showing 5,067 regions with increased chromatin accessibility (left panel) or 4,577 regions with decreased chromatin accessibility (right panel) in representative individuals. Accessibility determined by differential accessibility analysis with R package DiffBind using a log_2_ fold change cutoff of >|0.58| and a p-value <0.05 (E) Genomic distributions of each ATAC peakset, regions with increased or decreased accessibility, as annotated by HOMER annotatePeaks.pl (F) Motif enrichment with HOMER searching either regions with increased accessibility (upper panel) or decreased accessibility (lower panel), showing the putative transcription factors binding within these regions. (G) Changes in promoter and enhancer chromatin accessibility observed in *Prdm3/16ShhCre*^*Δ/Δ*^ epithelial cells in differentially expressed genes associated with AT2 cell maturation.

Since morphological studies were consistent with pulmonary immaturity, we used correlation analysis to estimate the overall degree of immaturity within each cell lineage. We generated epithelial cell type specific gene expression profiles from pseudo-bulk analyses of single cell RNA profiles from WT and *Prdm3/16*^*ShhCreΔ/Δ*^ animals and compared them with pseudo-bulk analysis of single cell RNA expression profiles from an embryonic mouse lung developmental time course.^44^ RNA profiles from WT AT1, AT2, and AT1/AT2 cells were all highly correlated with time-matched cell-specific signatures from AT1^Zepp^, AT2^Zepp^, and AT1/AT2^Zepp^. RNA profiles from *Prdm3/16*^*ShhCreΔ/Δ*^ AT2 cells were most correlated with those from E12.5 AT2^Zepp^, demonstrating that lack of PRDM3/16 was sufficient to prevent progression of cell differentiation within the AT2 lineage (Figure 2F). AT2 and AT1/AT2 cells consistently were more similar to immature rather than mature alveolar cells. RNA profiles from AT1 cells from *Prdm3/16*^*ShhCreΔ/Δ*^ pups were most concordant with E17.5 AT1^Zepp^, though less correlated than in WT controls, confirming that AT1 gene expression was only mildly altered by absence of PRDM3/16 (Figure 2F).

Strikingly, PRDM3/16 deletion caused E18.5 AT1/AT2 cells to most closely resemble the immature E12.5 and E15.5 SOX9 progenitor cells. These cells also continued to express low levels of *Sox9*. The immaturity of the *Prdm3/16*^*ShhCreΔ/Δ*^ AT2, AT1, and AT1/AT2 cells is also indicated by changes in gene expression shared between the cell types. We observed increased expression of genes previously reported^46^ to be downregulated during maturation of Sox9 progenitors including *Igfbp5, Pp1r14b, Peg3, Igf2, Cmas, Rbp1*, and *Tmsb10*, with concordant decreased expression of genes normally upregulated during progressive maturation including *Gde1, Matn4, Muc1, Nkd1, Mt1, Tspo, Ctsh, Brd7, Atp1b1, Tmbim6* and *Scd2*. Although the numbers of Sox9 high progenitors are few at this stage of development, the *Prdm3/16*^*ShhCreΔ/Δ*^ population expressed higher levels of markers associated with proliferation (Figure S3F). Together, these observations imply a failure of distal progression within the Sox9 progenitor population during later stages of embryonic lung development, emphasizing the role of PRDM3/16 in driving cell fate acquisition and maturation.

Gene expression changes observed in the single cell analysis were reflected in bulk RNA obtained from EpCAM^+^ MACS-sorted pulmonary epithelial cells from E17.5 *Prdm3/16*^*ShhCreΔ/Δ*^ and control lungs (Figures 3 and S4A, Table S2). Consistent with the scRNA and immunofluorescence data, genes associated with surfactant biosynthesis (*Sftpb, Sftpc*, and *Sftpd)*, genes associated with surfactant homeostasis (*Ctsh*) and those associated with lipid metabolism (*Lpcat1, Abca3, Fasn, Scd1)* were significantly decreased in bulk epithelial RNA (Figure 3B, C). Several genes not found to be differentially expressed in the single cell data but known to be required for normal lung development and AT2 cell maturation were differentially expressed in the bulk RNA analysis including *Fgfr2* (−1.73),^47,48^ *Cebpa* (−1.83),^49^ *Etv5* (−2.82),^50,51^ and *Wnt5a* (2.1)^52^(Figure 3A). RNAs associated with the main AT1 cell program were increased, including *Ager, Rtkn2, Lima1, Tead4, Cttnbp2*, and *Mslnl*, consistent with the increased number of AT1 cells (Figure 3A, C); however, a number of genes associated with AT1 cells were downregulated in both the bulk and single cell data including *Fbln5, Bmp4, Slco3a1, Samhd1, Slc4a5*, and *Gnb4*. Taken together, analysis of both single cell and bulk RNA gene expression indicates cellular immaturity across the late embryonic lung epithelium, especially in AT2-like cells, in the absence of PRDM3 and PRDM16.

### Chromatin landscape changes after loss of PRDM3/16

Because PRDM3 and 16 function as histone methyltransferases, we hypothesized that the differential gene expression and cellular immaturity found in *Prdm3/16*^*ShhCreΔ/Δ*^ fetuses was related to changes in chromatin organization. We used ATAC-seq to profile EpCAM^+^ sorted epithelial cells isolated from E17.5 control and *Prdm3/16*^*ShhCreΔ/Δ*^ lungs. Differential accessibility at 32,886 sites was identified (log_2_FoldChange > |0.58|, p-value <0.05), the majority of which annotated to intergenic or intronic regions (Figure 3D, E) at putative enhancer sites. Fewer differentially accessible sites were annotated to promoter regions (Figure 3E). Peaks with increased accessibility after deletion of *Prdm3/16* were enriched in transcription factor binding sites, including FOXA1/A2, NKX2-1, and TEAD, while peaks that showed decreased accessibility were enriched in the predicted transcription factor binding sites for NKX2-1, FOS/JUN, CEBP, and TEAD (Figure 3F). Multiple changes in chromatin accessibility were adjacent to genes associated with functional enrichment annotations that were overrepresented in bulk and scRNA data from *Prdm3/16*^*ShhCreΔ/Δ*^ animals (Figure S4C); of the genes differentially expressed in both the bulk and single cell data, 72% were associated with nearby changes in chromatin accessibility (Table S3). Notably, changes in accessible sites varied across a single genetic region with changes to both more open and more closed configurations when comparing WT and *Prdm3/16*^*ShhCreΔ/Δ*^ epithelium. Closure of the chromatin near transcriptional start sites (TSS) was observed in a subset of AT2 genes including *Cftr, Chia1, Fabp12, Lamp3, Napsa, Sftpd, Spink5, C3, Scd1, and Retnla*. Chromatin accessibility was also altered near *Cebpa, Elf5, Fabp5, Slc34a2, Pi4k2b, Insig1, Lpin2, Abca3, Fgfr2*, and *Atp6v1c2* (Figure 3G, Table S3). Increased chromatin accessibility was observed near the AT1 marker genes *Ager* and *Slco2a1*, consistent with increased expression at the RNA level. Taken together, our results suggest PRDM3 and PRDM16 activate and repress alveolar epithelial gene expression at least in part by modulating the accessibility of promoter and enhancer regions associated with the binding of lineage defining transcription factors including NKX2-1 and other co-factors such as CEBPA and TEAD.

### PRDM16, PRDM3, and NKX2-1 share DNA binding sites

Since the AT2 signature genes whose expression was decreased in the *Prdm3/16*^*ShhCreΔ/Δ*^ mice were consistent with those regulated by NKX2-1,^8,12^ and NKX2-1 binding sites were enriched at ATAC-seq peaks which were differentially accessible after *Prdm3* and *Prdm16* deletion, we hypothesized that PRDM3/16 functioned in alveolar epithelial cells via interaction with NKX2-1. We tested whether PRDM16, NKX2-1, and PRDM3 bound to shared genomic sites using CUT&RUN. Antibodies against PRDM16, NKX2-1, and PRDM3 were used to identify chromatin binding sites in bulk E17.5 EpCAM^+^ sorted epithelial cells. PRDM16 bound to 35,847 unique genomic regions, 22.5% near TSS (Figure 4A, B, Table S4). PRDM3 showed a similar distribution throughout the genome (Figure S5, Table S5). NKX2-1 bound to 52,716 unique genomic regions, 8,004 (∼18.5%) of which were located near the promoter-TSS regions (Table S6). Remarkably, PRDM16 and NKX2-1 bound together at ∼40% of the identified peaks and PRDM16, NKX2-1, and PRDM3 coordinately bound at ∼20% of the identified peaks. Motif enrichment of sequences in the shared binding regions included predicted binding sites for TEAD, FOXA, and NKX2-1 (Figure 4C, Figure S5), sites also enriched within the identified ATAC-seq peaks. The enrichment of TEAD binding sites within PRDM16 peaks was previously also observed in embryonic heart.^38^ Consistent with other PRDM16 Chip-seq data sets, the previously identified consensus binding sequence for PRDM16 was not identified within PRDM16, PRDM3, and NKX2-1 bound peaks.^32,38,53,54^

**Figure 4.**
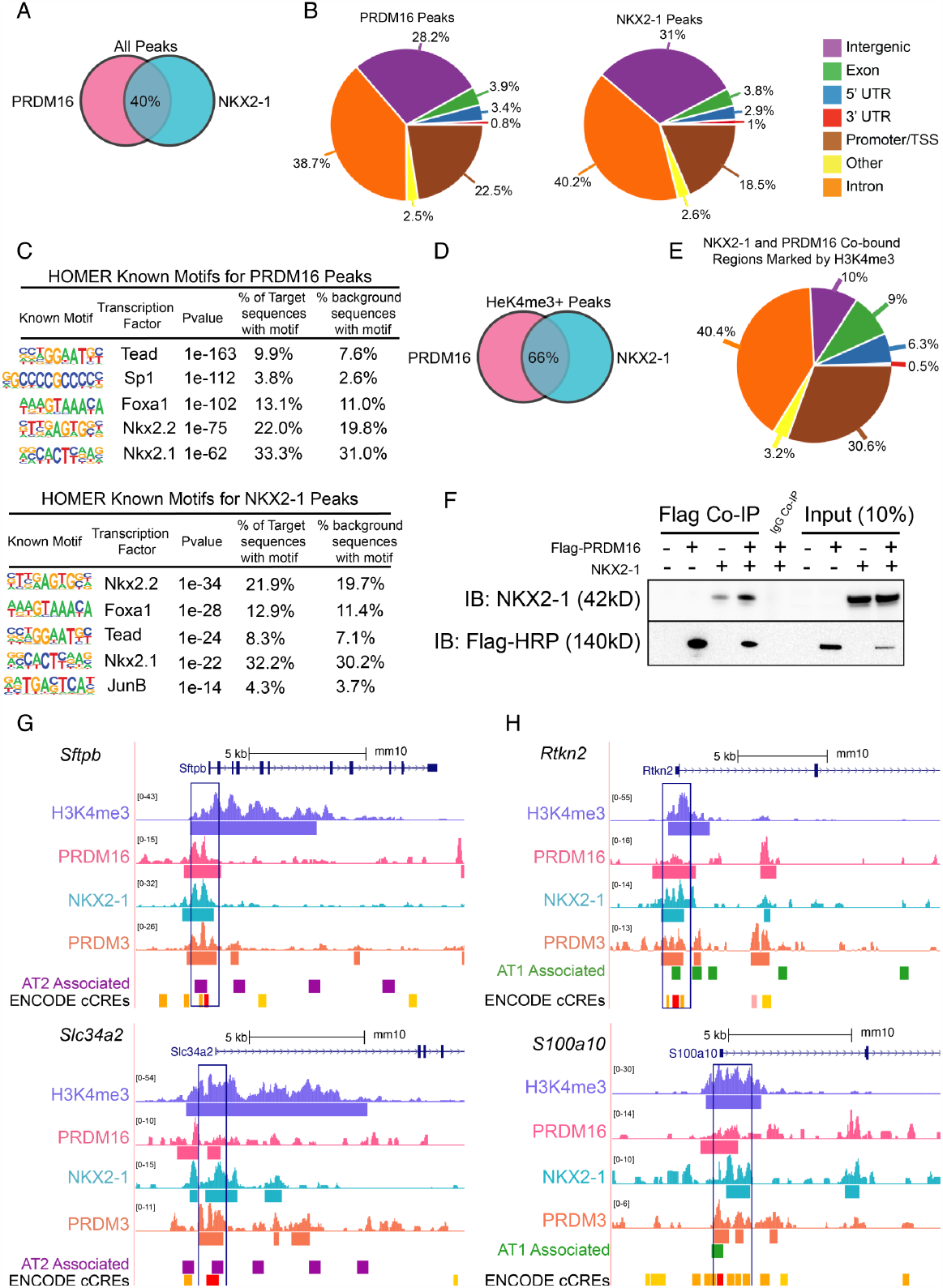
PRDM16, NKX2-1, and PRDM3 bind to shared sites throughout the genome and at promoters. (A) Venn Diagram shows the average overlap of binding sites between PRDM16 CUT&RUN and NKX2-1 CUT&RUN observed in two experiments. (B) Genomic distributions of all called peaks from a representative PRDM16 binding experiment (left panel) and a representative NKX2-1 binding experiment (right panel). (C) HOMER motif enrichment for all called peaks across the genome of a representative PRDM16 CUT&RUN (upper panel) or NKX2-1 CUT&RUN (lower panel) experiments. (D) Venn diagram of the overlap of H3K4me3 marked peaks between PRDM16 and NKX2-1 CUT&RUN. (E) Genomic distributions of the overlap peaks bound by both PRDM16 and NKX2-1 that are marked by H3K4me3. (F) Immunoprecipitation showing co-binding of FLAG-tagged PRDM16 and NKX2-1 after co-transfection in HEK293T cells. (G and H) CUT&RUN analysis of selected genes is visualized with the UCSC Genome Browser for H3K4me3, PRDM16, PRDM3, and NKX2-1. AT1 cell and AT2 cell associated peaks from published data set (Little, et.al.,(12)). ENCODE cCRE peaks are annotated from the ENCODE database of cis-regulatory elements. Binding is seen in AT2 cell-associated genes (G) and AT1 cell-associated genes (H).

We used CUT&RUN to identify regions bearing the chromatin mark H3K4me3 to identify regions of active transcription, and compared these regions to those bound with PRMD16, PRDM3, and NKX2-1. Analyzing only areas marked by H3K4me3, PRDM16 and NKX2-1 co-localized at 66% of peaks with 30.6% of these sites being located near the transcriptional start site (Figure 4D, E). Chromatin binding sites in active regions shared by NKX2-1, PRDM16, and PRDM3 were identified near AT2 cell signature genes including *Sftpc, Sftpb, Cebpa, Abca3, Slc34a2, Scd1*, and *Lamp3*, genes previously shown to be bound and activated by NKX2-1 (Figure 4G, Figure S5D, Table S7).^12,21,55^ Co-binding of PRDM16 and NKX2-1 was also observed near AT1 cell-associated genes, for example, *Bmp4, Pmp22, Sem3b, Slco2a1, S100a10, Cldn18, Col4a2*, and *Fgfbp1 (*Figure 4H, Supplemental Figure 5D and Table S7). Since PRDM16 and NKX2-1 shared occupancy at regulatory regions of target genes, we performed co-immunoprecipitation assays to evaluate direct binding. NKX2-1 and PRDM16 co-precipitated in co-transfected HEK293T cells, indicating a potential direct interaction between NKX2-1 and PRDM16 (Figure 4F).

We constructed a gene regulatory network (GRN) serially integrating present data from epithelial bulk RNA-seq, scRNA-seq, PRDM3, PRDM16, NXK2-1, and H3K4me3 CUT&RUN, and NKX2-1 ChIP-seq data from Little et al.^12^ First, we defined genes for the transcriptional regulatory network (TRN) input based on increased expression following PRDM3/16 deletion in AT1 (AT1 up regulated network) or decreased expression following PRDM3/16 deletion in AT1 (AT1 down regulated network) or AT2 cells (AT2 network). We filtered these gene sets for those genes with neighboring chromatin regions containing PRDM3, PRDM16 and NKX2-1 binding sites by CUT&RUN or in published NKX2-1 ChIP-seq data and H3K4me3-marked active chromatin. The identified genes were used as input for IPA analysis to predict key upstream regulators of the AT1 and AT2 networks (Figure 5, Figure S6).

Within the AT2 network CEBPA, ETV5, FGFR2, SREBF1, and STAT3 were predicted to have reduced activity following deletion of *Prdm3/16*, while CBX5, ID3, and FOXA1 were predicted as activating factors in the NKX2-1/PRDM3/16 centered network (Figure 5A). Other cofactors such as CBX5, ID3, and FOXA1/A2 were also predicted to be involved. The synergistic roles of CEBPA, ETV5, SREBF1, FOXA2, and GATA6 with NKX2-1 in AT2 cell maturation, differentiation and surfactant homeostasis were previously demonstrated.^7,21,49,56,57^ Co-factors active after PRDM3/16 deletion include CBX5, ID3, and FOXA1, were predicted to regulate the “negative regulation of transcription by RNA polymerase II (GO:0000122)” and “negative regulation of cell differentiation (GO:0045596)”. The promoter sequences of two genes expressed in AT2 cells, *Sftpb* and *Abca3* were scanned for binding motifs from the JASPAR and cis-BP motif databases.^58,59^ Binding motifs for CEBPA, FOXA1/A2, SREBF1, and STAT3 were identified in addition to NKX2-1 further confirming the network of predicted cofactors (Figure 5B). Within the *Sftpb* and *Abca3* promoters, the predicted cis-BP PRDM16 binding site was observed. This site was only identified in 15% of all the PRDM16 CUT&RUN peaks. Notably, within the *Abca3* promoter, the predicted cis-BP PRDM16 site was not directly associated with the identified CUT&RUN peaks suggesting that this sequence may not be the only mechanism through which PRDM16 regulates a genomic region. These analyses predict that PRDM3/16 function with NKX2-1 to activate gene regulatory networks required for AT2 differentiation and surfactant homeostasis before birth.

**Figure 5:**
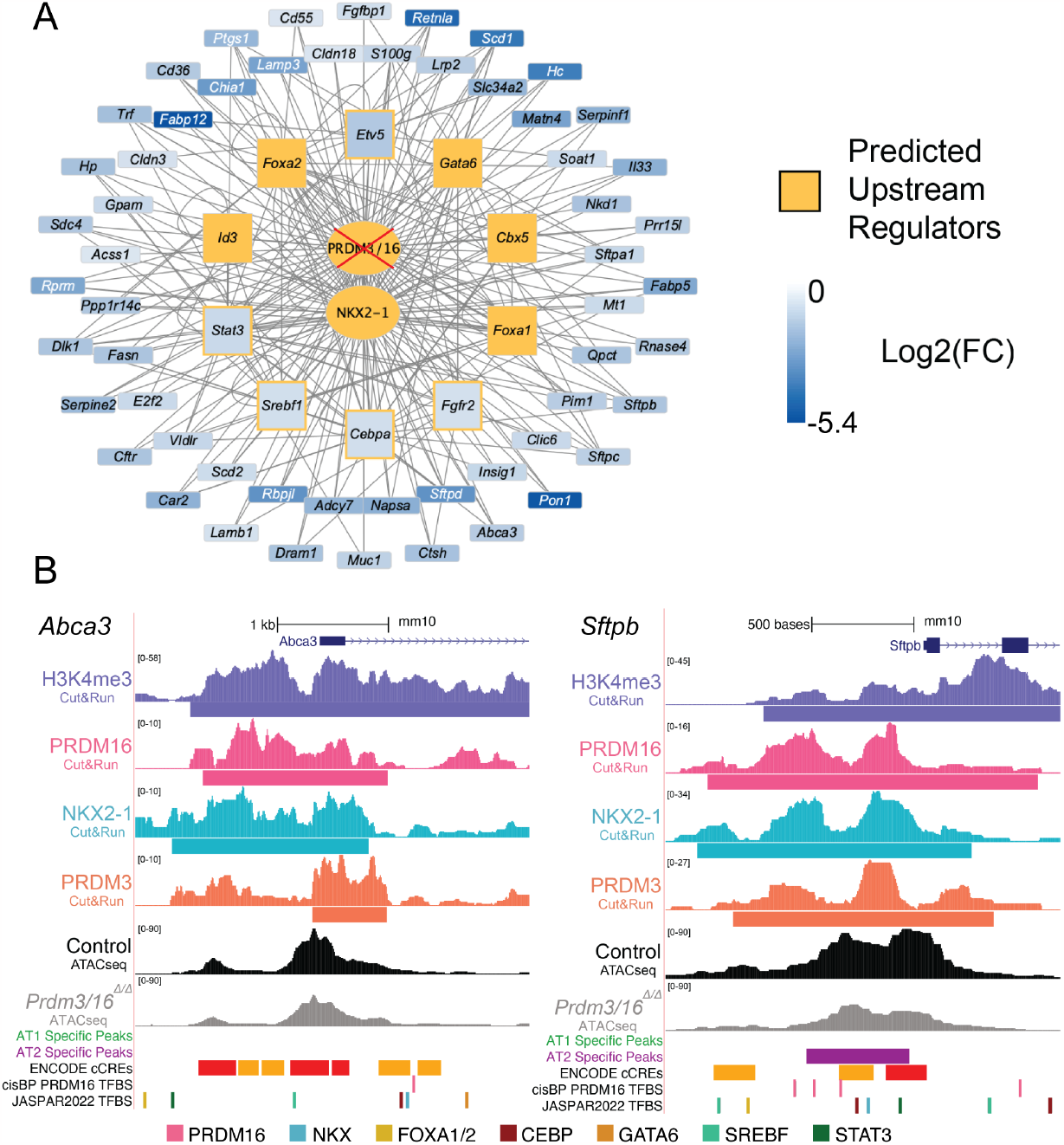
PRDM3/16 work alongside NKX2-1 to regulate gene networks critical for AT2 cell differentiation. (A) Gene regulatory network for AT2 cells with PRDM16 and NKX2-1 at the center. Genes were selected based on being downregulated in the bulk RNA-seq (Log_2_ fold change > |0.58|, p-val <0.05) and either differentially expressed in the single cell DEA or single cell gene expression (>15% expression in the AT2 control cell population and *Prdm3/16*^*ΔΔ*^ expression > Control expression). Those genes were used as input into IPA for a Regulatory Upstream Analysis. The resulting relationships were loaded into Cytoscape. IPA predicted previously identified transcription factors known to regulate key genes in AT2 cells. (B) Predicted transcription factor binding sites (TFBSs) of PRDM TFs and co-factors near the promoter regions of canonical AT2 genes. TFBSs visualization is based on the data of “JASPAR 2022 TFBS UCSC tracks” available in the UCSC genome browser. TFBSs with prediction scores >=400 were included and the locations of PRDM16, NKX (NKX2-1, NKX2-2), FOXA1/2, SREBF1/2, CEBP (CEBPA/D/G), GATA6, and STAT3 are indicated. The predicted PRDM16 binding motif from the CIS-BP database was also included using HOMER scanMotifGenomeWide.pl to scan reference sequences in the promoter regions of *Abca3* and *Sftpb*. Nearby TFBSs of the same TF family were merged. TFBSs were colored based on TF family.

The up-regulated AT1-associated gene network identified ARID1A, CTNNB1, HBEGF, and SMARCA4 as predicted co-factors active in the NKX2-1/PRDM centered network regulating AT1 cell gene expression (Figure S6A). NKX2-1, ARID1A, CTNNB1, and HBEGF are known to form complexes with SMARCA4.^60-63^ Both SMARCA4 and ARID1A are members of the SWI/SNF family involved in chromatin remodeling. ARID1A deficiency influences *Prdm16* expression in mouse embryo fibroblasts.^64,65^ The down-regulated AT1-associated network was comprised of the AT1 genes *Pdpn, Cldn18, Fbln5, Hs2st1, Phactr1*, and *Cxadr*, as well as genes normally expressed at high levels in AT2 cells, such as *Napsa, Cldn18, Sftpd, Rbpjl, Ctsh*, and *Muc1* with an enriched function on surfactant balance (GO:0043129) and alveolar development (GO:0048286). In addition to SMARCA4 and ARID1A; NKX2-1, GATA6, KRAS, STAT6, and RXRA were identified as key upstream regulators (Figure S6B). Taken together, these data implicate PRDM3/16 as direct co-regulators with NKX2-1, binding cooperatively across the genome to drive chromatin organization and define the AT1 and AT2 specification and differentiation programs during distal lung differentiation.

### Prdm3 and Prdm16 are required for AT2 lineage fidelity during lung development

The failure of AT2 cell differentiation in *Prdm3/16*^*ShhCreΔ/Δ*^ fetuses coupled with the observed early expansion of the AT1 expression domain and increase in AT1 cell numbers supported the hypothesis that PRDM3/16 loss in the AT2 lineage led to early transition of AT2-lineage cells to an AT1 cell fate. To directly test this hypothesis, we performed lineage analysis after AT2-lineage-specific PRDM3/16 deletion in *Prdm3/16*^*SftpcCreERT2*^; *Rosa26*^*lsl-tdTomato*^ animals. Recombination was induced by the administration of tamoxifen to the dams at E12.5 and E13.5, and the lungs were analyzed at E18.5 (Figure 6A). As previously reported,^43^ most WT cells marked by SFTPC^CreERT2^ early in lung morphogenesis express only AT2 cell markers including SFTPC at E18.5 (Figure 6B-I), with few cells expressing AT1 markers. In SftpcCreER^R26R-Tdt/*Prdm3/16*Δ/Δ^ fetuses (Figure 6J-Q), significantly more cells within the AT2 lineage (marked with TdTom) expressed HOPX alone (green arrowheads) or in combination with SFTPC (yellow arrowheads) while HOPX^-^ SFTPC^+^TdTom^+^ AT2 cells (white arrowheads) were decreased. These data demonstrate that PRDM3/16 function in the AT2 lineage is required for the fate acquisition and lineage fidelity during embryonic distal lung specification (Figure 6R-U).

**Figure 6:**
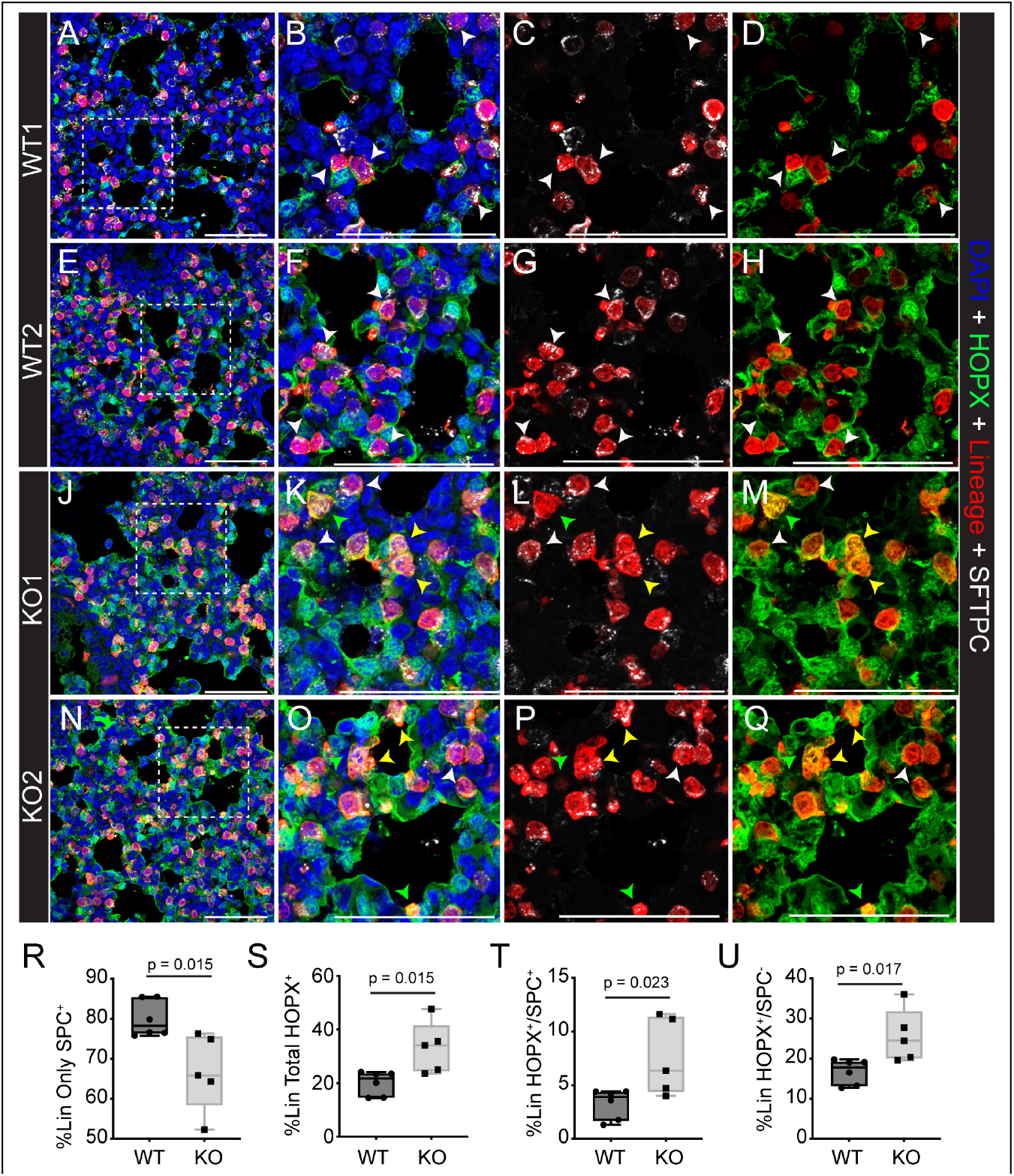
AT2-specific PRDM3/16 deletion leads to lineage infidelity during alveolar epithelial specification. Dams were treated with tamoxifen at E12.5 and E13.5 to generate SftpcCreER^R26R-Tdt^ (WT) and SftpcCreER^R26R-Tdt/*Prdm3/16*D/D^ (KO) fetuses which were harvested at E18.5 for lineage analysis. WT animals (A-H) demonstrated a majority of cells in the *Sftpc* lineage (Sftpc^Lineage^) were marked with only SFTPC protein by IHC (white arrowheads), while KO animals (J-Q) showed significant decreases in SFTPC-only cells ®, with corresponding increases in Sftpc^Lineage^ cells expressing HOPX (S), either with concomitant SFTPC expression (T) or without (U).

## Discussion

Respiratory function of the lung at birth requires interactions among a highly conserved network of signaling and transcription factors expressed by a diversity of cell types controlling lung morphogenesis and differentiation.^1,66^ Epithelial progenitors forming the embryonic respiratory tubules express the transcription factor NKX2-1 that distinguishes the initial foregut endoderm cells that are committed to pulmonary epithelial cell lineages.^1,4,43^ Thereafter, NKX2-1 is required for branching morphogenesis and differentiation of epithelial cells in conducting airways and alveoli, including specification of both surfactant-producing AT2 cells and thin, gas exchange promoting AT1 cells in the distal air sacks.^6,8,11,12^ Here we demonstrate that the histone lysine-methyl transferase proteins PRDM3 and 16 are required for AT1-AT2 cell fate choices. PRDM3/16 function together to promote AT2 fate acquisition and differentiation and participate in regulation of surfactant lipid and protein expression necessary for lung function at birth. They also are required for full expression of the AT1 program, leading to accumulation of partially differentiated AT1 cells in PRDM3/16 mutants. Transcriptomic and epigenomic studies demonstrated that PRDM3 and PRDM16 bind with NKX2-1 at predicted regulatory regions of their transcriptional target genes throughout the genome; without PRDM3 and PRDM16, chromatin accessibility at NKX2-1 bound promoter and enhancer sites is altered in AT2 and AT1 cell-associated genes. Thus, the present findings identify PRDM3 and 16 as novel epigenetic modulators of alveolar cell fate acquisition and specification required for full function of lung lineage transcription factors.

Loss of both *Prdm3* and *Prdm16* in alveolar epithelium caused surfactant deficiency at birth, consistent with the finding that both AT2 cell identity and maturation were inhibited in mutant lungs. The marked effect of PRDM3 and PRDM16 on AT2 cell lineage specification and differentiation supports the likelihood that PRDM3 and PRDM16 function in concert with other AT2-specific transcriptional co-activators in addition to NKX2-1, a concept supported by the co-localization of predicted binding sites for other transcription factors identified by sequence analysis of the CUT & RUN experiments. For example, multi-omics and GRN inference analysis predicted that FOXA1/A2, FOS/JUN, ETV5, GATA6 and CEBPA are involved in this biological process, all of which are known to regulate a similar group of AT2 cell signature genes and to bind to regulatory regions of NKX2-1 target genes.^7,49,56,57,67^ Impaired commitment to the AT2 cell lineage was previously observed in animals deficient in epithelial FGFR2B signaling during the late pseudo glandular period.^47,48,68^ Chromatin accessibility at the *Fgfr2b* locus was altered in *Prdm3/16* deficient epithelial cells accompanied by a reduction in the expression of *Fgfr2b* and one of its known targets, *Etv5*, also known to be critical for AT2 cell specification.^57,68^ The similarity in phenotype between these studies targeting signal transduction and chromatin regulation is provocative, suggesting identification of the precise integration of intercellular signaling with activity of chromatin complexes in modulating activity of partner transcription factors is an area of high priority for future studies.

While PRDM3 and PRDM16 and NKX2-bound together at many AT1 cell associated genes, AT1 cell gene fate and gene expression were generally maintained after deletion of *Prdm3/16*, indicating that the gene networks required for AT1 cell lineage choice and gene expression were substantially intact. Therefore, we conclude that the AT1-centric network is likely maintained by regulators and co-activators which are distinct from that required for AT2 cell differentiation. IPA analysis based on single cell and bulk epithelial RNA from AT1 cells is distinct from that predicted for AT2 cells, suggesting important roles for PRDM3/16 and NKX2-1 and their interactions with chromatin modulators and co-activators including SMARCA4, ARID1A, CITED2, and HIF1α. CITED2 is required for AT1 cell specification, activates SMAD2/3, directly interacts with HIF1a, activates the *Cebpα* gene promoter in late gestation and is required for AT2 cell differentiation.^69^ ARID1A and SMARCA4 function together as members of the SWI/SNF chromatin regulator complex, suggesting that the interaction of PRDM/NKX2-1 and the SWI/SNF complex may be vital for sustaining the epigenetic landscape essential for the AT1 lineage. It appears that loss of PRDM attenuates this complex interplay; we note a compensatory elevation in the activity of SMARCA4 and ARID1A in PRDM3/16 knockouts, presumably as a homeostatic response to preserve the integrity of the AT1 cellular environment. In combination, this leads to increased expression of AT1-specific genes and an expansion of the AT1 cell population, further underscoring the dynamic nature of the epigenetic regulation within AT1 cells. We speculate that such interplay of epigenetic factors likely adds a distinct layer of regulation across the period of lung epithelial specification.

Finally, it is intriguing to note that lineage-specific deletion of PRDM3/16 led to partial lineage infidelity in the AT2 lineage during distal lung specification. Extensive recent literature has assessed the relationship between AT1 and AT2 lineages during alveolarization^43,47,70,71^ with some data supporting a model of early lineage specification by E14.5^43^ and other data suggesting a population of AT1/AT2 cells retained in the late prenatal lung.^71^ In our study, loss of PRDM3/16 led to a decrease in the percentage of AT2 cells derived from early SFTPC-expressing epithelium at E12.5 from 80% to 60%, with concomitant increases in both HOPX^+^/SFTPC^-^ AT1 cells and HOPX^+^/SFTPC^+^ AT1/AT2 cells. Combined with data suggesting that AT1/AT2 cells in *Prdm3/16*^*ShhCreΔ/Δ*^ animals were most similar to earlier progenitors, these data suggest that one key role of PRDM3/16 is to refine the lineage potential of distal lung epithelium. Therefore, functional chromatin remodeling via PRDM3/16 leads to restricted, refined lineages with appropriate quorum and fate identity, while loss of chromatin architecture prevents appropriate fate specification and leads to retention of more plastic, less mature, and less differentiated cells. Control of cellular plasticity is critical to both lung development and regeneration following lung injury, and we speculate that definition of the epigenetic regulators, like PRDM3/16, which regulate plasticity versus differentiation may open novel therapeutic avenues for lung disease across the lifespan.

## Acknowledgements

The authors thank members of the CCHMC Single Cell Genomics Facility (RRID:SCR-022653) and the CCHMC Bio-imaging and Analysis Facility (RRID:SCR-022628) for their advice and support for sample generation and analysis. We also thank Dr. Patrick Seale for generously providing the Rabbit anti-PRDM16 antibody. This research was supported by grants from the National Key R&D Program of China (2022YFA0806200), NSFC (82200098), and the Fundamental Research Funds for the Central Universities (SCU2022D006) to Hua He and NHLBI R01HL164414 to Jeffrey Whitsett and William Zacharias.

## Author Contributions

HH and SMB designed and conducted experiments and prepared the manuscript. AKD and JS performed ATAC, Bulk RNA, and CUT & RUN bioinformatic analyses and figure preparation, CLN performed EM analysis, AS immunoprecipitation studies, MG and SZ performed single cell analysis and figure preparation, YX created the GRN, WJZ prepared the manuscript. MK and SG provided *Prdm3 flox* mice, JAW designed experiments and prepared the manuscript.

## Competing interests

The Authors declare that they have no competing interests for the current work, including patents, financial holdings, advisory positions, or other interests.

## Materials and Methods

### Animals

All animals were housed in specific pathogen-free facilities under the IACUC protocols approved by Cincinnati Children’s Hospital and West China Second University Hospital of Sichuan University. Mice were bred on a mixed background and housed with food and water ad libitum. *The Prdm16*^*flox/flox tm1*.*1Brsp*^/J(strain 024992, RRID:IMSR_JAX:024992),^42^ *Shh*^*tm1(EGFP/cre)Cjt*^/J (strain 005622, RRID:IMSR_JAX:005622),^86^ *Gt(ROSA)26Sor*^*tm9(CAG-tdTomato)Hze*^/J (strain 007909, RRID:IMSR_JAX:007909),^87^ C57BL6/J (strain 000664, RRID:IMSR_JAX:000664) mice were acquired from Jackson Laboratories (Bar Harbor, ME). *Sftpc*^*CreERT2*^ mice were obtained from Dr. Harold Chapman.^73^ *Prdm3*^*flox/flox*^ mice were acquired from Dr. Kurokawa.^41^ For *ShhCre* breeding, *Prdm3*^*flox/flox*^*Prdm16*^*flox/+*^*Shh*^*Cre*^ males were crossed with *Prdm3*^*flox/flox*^*Prdm16*^*flox/flox*^ females to generate double knockout embryos, *Prdm3*^*flox/flox*^*Prdm16*^*flox/flox*^ littermates were used as controls. For the lineage tracing study, *Prdm3*^*flox/+*^*Prdm16*^*flox/+*^*Sftpc*^*CreERT2*^ males were interbred with *Prdm3*^*flox/flox*^*Prdm16*^*flox/flox*^*ROSA26*^*tdTomato/tdTomato*^ mice; Cre positive, double heterozygous littermates (*Prdm3*^*flox/+*^*Prdm16*^*flox/+*^*Sftpc*^*CreERT2*^*ROSA26*^*tdTomato/+*^) were used as controls. Animals were mated overnight, and the presence of a vaginal plug the next morning was defined as E0.5. To induce CreER activity, *Sftpc*^*CreERT2+*^ pregnant dams were given two doses of tamoxifen (Aladdin Scientific, T137974) dissolved in corn oil (20 mg/ml, 200 mg/kg) at E12.5 and E13.5 via oral gavage.

### Histology and Immunofluorescence Analysis

Embryos were harvested and fixed in 4% paraformaldehyde (Electron Microscopy Sciences) in PBS at 4°C overnight. Tissues were rinsed in PBS and either dehydrated through graded ethanols and into xylene prior to paraffin embedding or equilibrated in 30% sucrose/PBS prior to being embedded in OCT for cryosections. 5 μm paraffin sections were stained with Hematoxylin and Eosin (H&E), and images acquired on a Nikon Eclipse Ti2 microscope. Immunofluorescence staining was performed on 5 μm sections that underwent heat mediated antigen retrieval in citrate buffer (pH 6.0).

Primary antibodies used include: Mouse anti-HOP (Santa Cruz Biotechnology sc-398703; 1:100), Guinea Pig anti-pro-SFTPC (In house; 1:500), Rabbit anti-ABCA3 (Seven Hills Bioreagents WRAB-70565; 1:100), Rabbit anti-NKX2.1 (Seven Hills Bioreagents WRAB-1231; 1:1000), Mouse anti-SOX2 clone E-4 (Santa Cruz Biotechnology SC-365823; 1:100), Rabbit anti-SOX9 (Millipore AB5535; 1:100), Guinea Pig anti-LPCAT1 (In house,1:300),^72^ Rabbit anti-Aquaporin5 (Abcam ab78486; 1:1200), Hamster anti-Podoplanin (DSHB 8.1.1: 1:200), Rat anti-Ager (R&D systems mab1179; 1:100 no antigen retrieval), Guinea Pig anti-Lamp3 (Synaptic Systems 391005; 1:500), Sheep anti-PRDM16 (R&D systems AF6295; 1:100), Goat anti-endomucin (R&D systems A45666: 1:1000), Rabbit anti-Pro SFTPC (Sigma AB3786; 1:400), and Goat anti-Tdtomato (Origene AB8181; 1:500).

Samples were incubated with primary antibodies overnight in a humid chamber at 4°C. After 3 times washing with PBS-TritonX100 (0.1%), sections were labeled with fluorophore-conjugated or biotinylated secondary antibodies. Secondary antibodies were all used at a dilution of 1:200 and include Thermo Fisher Scientific antibodies: Goat anti-mouse IgG1 Alexa Fluor 647 (A21240), Goat anti-Guinea Pig IgG Alexa Fluor 568 (A11075), Goat anti-Rabbit IgG Alexa Fluor 488 (A-11034), Goat anti-Rat IgG Alexa Fluor 647 (A21247), Goat anti-Mouse IgG1 Alexa Fluor 568 (A21124), Donkey anti-Rabbit IgG Alexa Fluor 647 (A31573), Goat anti-Hamster IgG Alexa Fluor-488 (A21110), Donkey anti-Goat 647 (A21447), and Jackson ImmunoResearch antibodies Biotin-conjugated Donkey anti-sheep (713-065-147), Alexa Fluor-488 Donkey anti-Guinea Pig IgG (706-545-148), Alexa Fluor-647 Donkey anti-rat IgG (712-605-153), Alexa Fluor-488 Donkey anti-Rabbit IgG (711-545-152), Alexa Fluor-594 Donkey anti Mouse IgG (715-585-151), and Alexa Fluor-647 Donkey anti Goat IgG (705-605-147). A Tyramide Biotin Signal Amplification (TSA) kit (Akoya/Perkin Elmer) was used for PRDM16 signal amplification in combination with Streptavidin, Alexa Fluor-488 (Life Technologies 532354; 1:200). SCD1 immunohistochemical staining was performed using Rabbit anti-SCD1 (Abcam 236868; 1:200 with TRIS EDTA pH9.0 antigen retrieval), Goat Biotinylated anti-Rabbit IgG (Vector Labs BA-1000), the Vectastain Elite ABC Peroxidase kit (Vector Laboratories: PK-6100) and nuclear fast red post staining. Sections were imaged on Nikon A1R, Leica Stellaris 5, or Olympus IxploreSpin confocal systems with identical laser exposure between groups.

For AT1 and AT2 cell quantification, six 60x confocal images taken from at least three different lobes at the distal margins were used. Z-stack images comprised of 18 positions were denoised using Nikon Elements software. To reduce the background contributed by red blood cells, the signal from the TRITC channel was subtracted from the FITC channel to create a new FITC-TRITC channel. Files were analyzed in IMARIS. Spot counting was performed with a nucleus size of 5 mm for DAPI and NKX2.1^+^ counting. A 6 mm area size was used for HOPX staining. Automatically counted DAPI^+^, NKX2.1^+,^ and HOPX^+^ spots were manually evaluated, and corrections were made as necessary. The presence of SFTPC was manually scored. The number of AT2 cells was determined by filtering SFTPC+ cells with the shortest distance to DAPI and NKX2.1 and the number of AT1 cells by filtering HOPX+ cells with the shortest distance to DAPI and NKX2.1. AT1/AT2 cells were DAPI^+^/NKX2.1^+^/SFTPC^+^/HOPX^+^ as defined by IMARIS and manually confirmed/corrected. For the lineage tracing study, six to eight 60x confocal images were taken from at least three different lobes. SFTPC, tdTomato, and nuclear HOPX signals were counted manually using the “Cell Counter” plug-in in FIJI software.

### Single cell RNA sequencing

E18.5 lungs were dissected, and extrapulmonary tissue was removed from embryos isolated from two litters. Samples were stored on ice in HypoThermosol FRS preservative solution (Stem Cell Technologies, 07935) until genotypes were determined. For each sample, two lungs of the same genotype were minced together using a razor blade for 2 minutes to create a fine paste. 25 mg of minced tissue was digested on ice for 4 minutes in 1 ml of Collagenase/Elastase/Dispase enzyme mix (6 mg/ml Collagenase A (Sigma, 10103578001), 4.3 units/ml Elastase (Worthington, LS002292), 9 units/ml Dispase (Worthington, LS02100), 5mM CaCl2, 125 Units/ml DNAse (Applichem, A3778), in Dulbecco’s phosphate buffered saline (Thermo Fisher, 14190144). The tissue was returned to a sterile petri dish and minced for 2 minutes with a razor blade. The suspension was pipetted back into an Eppendorf tube on ice; the dish was rinsed with an additional 0.5 ml of enzyme mix. The mixture was triturated and shaken for an additional 2 minutes for a total digestion time of ∼8 minutes. The remaining tissue chunks were allowed to settle, and ∼1200 ml of the supernatant passed through a 30 mm filter (Miltenyi, 130-098-458) on a 15 ml conical. The filter was rinsed with 5 ml ice-cold PBS/BSA 0.04%. And the suspension held on ice. While on ice, the original tube with remaining chunks of tissue was further digested with 1 ml of enzyme mix containing (10 mg/ml Bacillus Licheniformis (Sigma, P5380), 0.5mM EDTA, in Dulbecco’s Phosphate Buffered Saline (Thermo Fisher, 14190144) with periodic trituration for an additional 10 minutes prior to passing through the retained 30 mm filter and additional rinsing of the filter with 7 ml PBS/BSA (0.04%). Cells were pelleted by centrifugation at 300xg for 5 minutes at 4°C, all but ∼100 ml of the supernatant aspirated, and cells were resuspended in 900 ml Red Blood Cell Lysis buffer (Sigma, R7757) and incubated on ice for 2 minutes. 12 ml PBS/BSA was added to the cells prior to centrifugation 200Xg for 5 minutes at 4°C. The supernatant was removed, and cells were resuspended in 1ml PBS/BSA. A step-by-step protocol can be found at https://www.protocols.io/view/adult-mouse-lung-cell-dissociation-on-ice-ymgfu3w. Cells were counted using a hemocytometer in the presence of trypan blue, and the concentration was adjusted to 1000 cells/ml prior to separation using the Chromium 10X Platform. Cell separation, cDNA synthesis, and library preparations were performed by the Cincinnati Children’s Hospital Gene Expression Core, and the libraries generated were sequenced by the Cincinnati Children’s DNA Core.

### Transmission electron microscope

Lungs were fixed in EM fixation buffer (2% glutaraldehyde, 2% paraformaldehyde (Electron Microscopy Sciences), 0.1% calcium chloride in 0.1M sodium cacodylate buffer pH 7.2) at 4°C overnight. Lung lobes were cut into 1-2 mm blocks and processed as previously described.^88^ Images were acquired by an H7650 transmission electron microscope (Hitachi).

### RNAscope

RNAscope was performed on paraffin-embedded 5 mm tissue sections from embryos of the indicated ages using the RNAscope Multiplex Fluorescent assay v2 according to the manufacturer’s instructions (Advanced Cell Diagnostics). Antigen retrieval was performed in a boiling water bath for 15 minutes. Sections were digested with Protease Plus for 15 minutes at 40°C. Probes used included Mm-Prdm16-C3 584281-C3 lot 22118C (1:50) and Mm-Mecom 432231 lot 22067A. Opals were assigned as noted: *Prdm3* (Opal 570, Akoya OP-001003 Lot 20212821, 1:500 incubated 45 minutes), and *Prdm16* (Opal 690, Akoya OP-001006 Lot 20211911, 1:500 incubated 30 minutes). Sections were post-stained with DAPI and imaged on a Nikon A1R-inverted confocal microscope.

### Isolation of bulk epithelial cells with magnetic-activated cell sorting

Lungs from E17.5 mice were minced into 1 mm^3^ pieces and incubated in RPMI media (Gibco, 11875-093) containing Liberase TM (Roche, 540111901, 50 μg/ml) and DNAse I (MilliporeSigma, 100 μg/ml) at 37 °C for 30 minutes. Tissues were transferred to C-tubes (Miltenyi) and dissociated with the GentleMACS cell dissociator. Single cells were subjected to RBC lysis buffer for 2 minutes on ice (eBiosciences, 00-4333-57) and incubated with FcR blocking reagent (Miltenyi, 130-092-575) on ice for 10 minutes, followed by incubation with CD326 anti-Mouse-EpCAM microbeads (Miltenyi, 130-105-958) at 4 °C for 15 minutes. Cells were then passed through LS columns for positive selection and collected in MACS buffer (Miltenyi). Cells were quantified manually under a hemocytometer with trypan blue before downstream experiments.

### RNA-seq, ATAC-seq, and CUT&RUN library construction

Following the EpCAM MACS sorting, 200,000 epithelial cells were aliquoted for RNA isolation with Rneasy Micro Plus kit (Qiagen, 74034) according to the manufacturer’s protocol. RNA samples were sent to Genewiz for polyA enrichment, library construction, and sequencing. ATAC-seq libraries were constructed as previously described.^89^ Briefly, 50,000 EpCAM^+^ epithelial cells isolated from control or *Prdm3/16*^*ShhCreΔ/Δ*^ E17.5 lungs were aliquoted, and the nuclei were isolated using lysis buffer (10 mM Tris-HCl, pH 7.4, 10 mM NaCl, 3 mM MgCl_2_, 0.1% IGEPAL CA-630). The nuclei were subjected to a transposition reaction with Illumina’s Tagment DNA TDE1 Enzyme and Buffer Kit (Illumina, 20034197) at 37°C for 30 minutes. Transposed DNA fragments were quantified by qPCR and amplified with barcoded primers using the NEBNext High-Fidelity 2X PCR master mix. The libraries were sent for sequencing by Genewiz and were sequenced on Hiseq4000 with PE150 mode for at least 50 million reads per sample.

The CUT&RUN experiments were performed using the Cut and Run Assay Kit (Cell Signaling Technology, 86652S). EpCAM^+^ epithelial cells from c57Bl/6 E17.5 mouse lungs were isolated and ∼100,000 cells were bound to concanavalin A beads and then incubated overnight with antibodies for PRDM16 (Sheep anti-PRDM16 R&D AF6295; RRID:AB_10717965; 1:50 or Rabbit anti-PRDM16 1:100 kindly provided by Dr. Patrick Seale ^34^), Rabbit anti-NKX2-1 (Seven Hills Bioreagents WRAB-1231; RRID:AB_2832953, 1:100), Rabbit anti-PRDM3 (Cell Signaling Technology 2593; RRID:AB-2184098, 1:50), Rabbit anti-H3K4me3 (Cell Signaling Technology 9751; RRID:AB_2616028,1:50), or rabbit IgG or Sheep IgG for each reaction. The cells were incubated with pAG-Mnase to facilitate antibody-guided DNA digestion. DNA was purified by phenol-chloroform-isoamyl alcohol (Thermo Fisher Scientific, 15593-031) extraction using Phasemaker tubes (Thermo Fisher Scientific, A33248) and ethanol precipitated. DNA was quantified using a Qubit and the dsDNA Quantification Assay Kit (Thermo Fisher, Q32850) or the Pico-green Assay Kit (Thermo Fisher, P7589). Libraries were constructed using the DNA Library Prep Kit for Illumina Systems (Cell Signaling Technology, 56795S) and Multiplex-oligos for Illumina Systems (Cell Signaling Technology, 47538S). Libraries were sequenced by Genewiz on a Hiseq4000 with PE150 mode for at least 15 million reads per sample.

### Co-immunoprecipitation

HEK293T cells were plated at a density of 2.8×10^5^ cells in 6-well plates and transfected with MSCV-flag-PRDM16 (Addgene, 15504; RRID:MSCV PRDM16) and/or pCDNA3-NKX2-1^74^ using Lipofectamine 3000 (Thermo Fisher Scientific, L3000075). Cells were lysed 48 hours post-transfection with Dynabeads Co-Immunoprecipitation kit lysis buffer (Thermo Fisher Scientific, 14321D) supplemented with 150mM NaCl, 2mM MgCl_2,_ and Complete, Mini Protease Inhibitor Cocktail (Roche, 11836153001) and PhosSTOP (Roche, 4906837001). Clarified lysates (∼800 μg) were pre-cleared with Protein A/G PLUS-agarose (Santa Cruz Technology, sc-2003) and non-specific IgG for 45 minutes, before incubating overnight with Ezview Red Anti-FLAG M2 Affinity Gel (Sigma-Aldrich, F2426; RRID:AB_2616449) at 4°C. Ezview Red Protein G Affinity Gel (Sigma-Aldrich, E3403) was used for negative control samples. Samples were washed with lysis buffer followed by 1X Tris-Buffered Saline buffer as per manufacturer’s protocol. Samples were eluted by boiling in Laemmli buffer with 5% β-Mercaptoethanol. Eluted proteins were resolved be SDS-page western blotting using 4–12% NuPAGE Bis-Tris gels (Thermo Fisher Scientific, NP0321BOX). Blots were incubated with Rabbit mAb anti-DYKDDDK Tag (HRP Conjugated) (Cell Signaling Technology, 86861, 1:1000, RRID:AB_2800094) or NKX2-1 antibody (Seven Hills Bioreagents, WRAB-1231, 1:1000) followed by incubation with TrueBlot HRP-Conjugated secondary antibody (Rockland Immunochemicals Inc. 1:1000, 18-8816-33, RRID:AB_2610848). Chemiluminescence detection was performed using Luminata Forte Western HRP substrate (MilliporeSigma, WBLUF0100) and a ChemiDoc Touch Imaging System (BioRad). Results shown are representative from at least three independent experiments performed.

### Analysis of 10x single cell RNA-seq of control and Prdm3/16^ShhCreΔ/Δ^ mouse lung

Sequencing reads from each single cell RNA-seq (scRNA-seq) sample were preprocessed, aligned, and quantified using the CellRanger pipeline (version 6.1.2, 10x Genomics) with mm10 mouse reference genome (refdata-gex-mm10-2020-A, 10x Genomics). For control and *Prdm3/16*^*ShhCreΔ/Δ*^ samples, the following cell filtering quality control (QC) steps were performed. EmptyDrops^75^ was used to identify cells (>100 unique molecular identifiers [UMIs] and false discovery rate <0.01) which deviated from empty droplets. Cells with 1,000-8,000 expressed (UMI > 0) genes, less than 100,000 UMIs, and less than 10% of UMIs mapped to mitochondrial genes were kept. Doublet cells in each sample were predicted and removed using Scrublet.^76^ In total, 18,084 cells from the four scRNA-seq samples passed the QC filters and were used for downstream analysis. Potential ambient RNA contamination in gene expression of each sample was corrected using SoupX^77^ with automatically estimated contamination fraction rates. SoupX-corrected data from individual samples were integrated using Seurat (version 4.1.0)^78^ reciprocal principal component analysis (RPCA) pipeline with SCTransform normalization. Cell clusters were identified using the Leiden clustering algorithm.^90^ An automated cell type annotation was performed using SingleR^79^ using LungMAP mouse lung Cellref^45^ as the reference. Results from the clustering analysis and the cell type annotation analysis were combined, and the final cell types were defined at the cluster level. Four clusters with co-expression of red blood cell genes (*Gypa, Alas2, and Bpgm*) and mesenchymal, epithelial, and endothelial cell markers were considered as contaminated clusters. Cell type annotations were validated using the expression of cell type marker genes from LungMAP CellCards and CellRef.^2,45^ Differential expression analysis was performed using the Seurat (v4.1.0) FindMarkers function with the Wilcoxon rank sum test. Differentially expressed genes were defined based on the following criteria, including adjusted *p*-value of Wilcoxon rank sum test <0.1, fold change ≥ 1.5, and expression percentage ≥20%.

For pseudo-bulk correlation analysis, gene expression matrix and cell type annotation of 10x single cell RNA-seq of mouse lung time course^44^ were downloaded from Gene Expression Omnibus using accession number GSE149563. Data from alveolar epithelial cell populations (AT1, AT2, and Epi progenitor cells) from E12.5, E15.5, and E17.5 were used for pseudo-bulk correlation analysis with the present scRNA-seq data of AT1, AT1/AT2, and AT2 cells from control and mutant mouse lungs at E18.5. Gene expression was normalized using Seurat LogNormalize function. Pseudo-bulk expression profiles for each cell population were calculated using Seurat AverageExpression function. Highly variable genes (HVGs) among pseudo-bulk profiles of each dataset were identified using Seurat FindVariableFeatures function. HVGs common in both datasets were used; then, the pseudo-bulk expression of those HVGs was z-score-scaled within each dataset. Pearson’s correlations of pseudo-bulk profiles were calculated using z-score-scaled expression of HVGs in the pseudo-bulk profiles.

### Bulk RNA-seq

Preprocessing of data from the Epcam^+^ sorted cells was done by using TrimGalore (0.4.2) and Cutadapt (1.8.1) to remove Illumina adapters. Quality was assessed with FastQC. Alignment was done with Bowtie (2.4.2) using the mm10 reference genome and the options ‘–end-to-end’ and ‘–sensitive.’ Low-quality alignments were removed with Samtools (1.10.0), and PCR duplicates were removed with Picard (2.18.22) MarkDuplicates. RNA-seq raw reads were aligned to the mm10 reference genome with Bowtie2. Raw count matrix was generated with HTSeq-count (2.0). DESeq2 (1.36.0) was used for differential analysis, with cutoffs of p<0.05 and log2 fold change > |0.58|. Heatmaps were generated with the pheatmap (1.0.12) R package, and the volcano plot was generated with EnhancedVolcano (1.14.0) R package. GO term enrichment was conducted using the lists of up and down differentially expressed genes as input on the ToppFun website. Dot plots were generated in R with ggplot2 (3.4.0). Representative pathways were selected based on both p-value and inclusion of classic AT1 or AT2 marker genes.

### Bulk ATAC-seq

Data files from Epcam^+^ sorted cells were processed following the standard ENCODE pipeline. Briefly, adapters were trimmed with Cutadapt (1.9.1) and aligned with Bowtie2 (2.2.6) to the mm10 reference genome. Mitochondrial low-quality reads (Samtools (1.7) view -F 1804), and duplicates (Picard 1.126) were filtered out. The IDR conservative peakset called by ENCODE pipeline defaults (MACS, p-value <0.01), along with the trimmed, sorted, deduplicated, no mitochondrial DNA bam file, were used for a DiffBind (v3) differential accessibility analysis, using full library normalization. A log_2_ fold change > |0.58| and p-value <0.05 were used to determine which regions showed increased or decreased accessibility under KO conditions. Statistically significant regions for both conditions were annotated with HOMER ‘annotatePeaks.pl’, and ‘findMotifsGenome.pl’ was used to determine enriched motifs.

### CUT&RUN

The CUT&RUNTools package was used for initial fastq file processing, with default settings for alignment, most notably the ‘–dovetail’ option in Bowtie2 (2.4.5). The MACS (2.1.4) package was used for peak calling with the following parameters: ‘-p 0.01 –keep-dup all’, using the -c option with an Igg control for each antibody; a p-value of 0.001 was used to call the H3K4me3 peaks. The narrow peak setting was used for all peak calls. Homer (4.11)^81^ ‘mergePeaks’ function was used to generate consensus peak sets across datasets. Homer (4.11) functions ‘annotatePeaks.pl’ and ‘findMotifs.pl’ were used to find the nearest genes and enriched motifs. The ‘bdgcmp’ function of MACS2 was used to normalize bedgraph files with fold enrichment.

### Genetic Regulatory Network Construction

The bulk RNA-seq, scRNA-seq; PRDM16, NKX2-1 and Hek4me3 CUT&RUN data, and the NKX2-1 ChIP-Seq data sets^12^ were used for network generation. Genes passing the following criteria were used for AT1 GRN construction: 1) gene expression increased in EpCAM+ sorted epithelial cells of *Prdm3/16*^*ShhCre*Δ/Δ^ vs. control (log_2_fold change >|0.58| p-value <0.05); 2) genes present in control AT1 cells (>15%) and percentage expression is higher in *Prdm3/16*^*ShhCre*Δ/Δ^ AT1 cells vs. control AT1 cells); and 3) genes with positive binding of PRDM3, PRDM16, NKX2-1 and H3K4me3 transcriptional activation marks based on the CUT&RUN data. In contrast, genes pass the following criteria were used for AT2 GRN construction: 1) gene expression decreased in EpCAM+ sorted epithelial cells of *Prdm3/16*^*ShhCre*Δ/Δ^ vs. control (log_2_fold change >|0.58| p-value <0.05); 2) genes present in control AT2 cells (>20%) and percentage expression was lower in *Prdm3/16*^*ShhCre*Δ/Δ^ AT2 cells vs. control AT2 cells); and 3) genes with positive binding of PRDM3 and16 and NKX2-1 based on the Cut & Run data. The resulting gene lists were subjected to Ingenuity Pathway Analysis (IPA) upstream regulator analysis. Activation and inhibition of key factors were determined by correlations between the observed expression changes of the target genes when compared to literature knowledge. Transcriptional Regulatory Networks (TRN) were generated to reveal the potential biological interrelationships among the key upstream regulators and their target genes. IPA was used to predict inhibited or activated upstream regulators based on the gene expression changes of the input genes.

### Data Accession

The Gene Expression Omnibus accession number associated with this data is GSE250366.

### Statistics

Data are presented as means ± SEM, and an unpaired 2-tailed t-test was performed to determine the significance between 2 groups. A p-value of less than 0.05 was considered significant. GraphPad Prism software was used for analysis and graph plotting.

## Key Resources Table

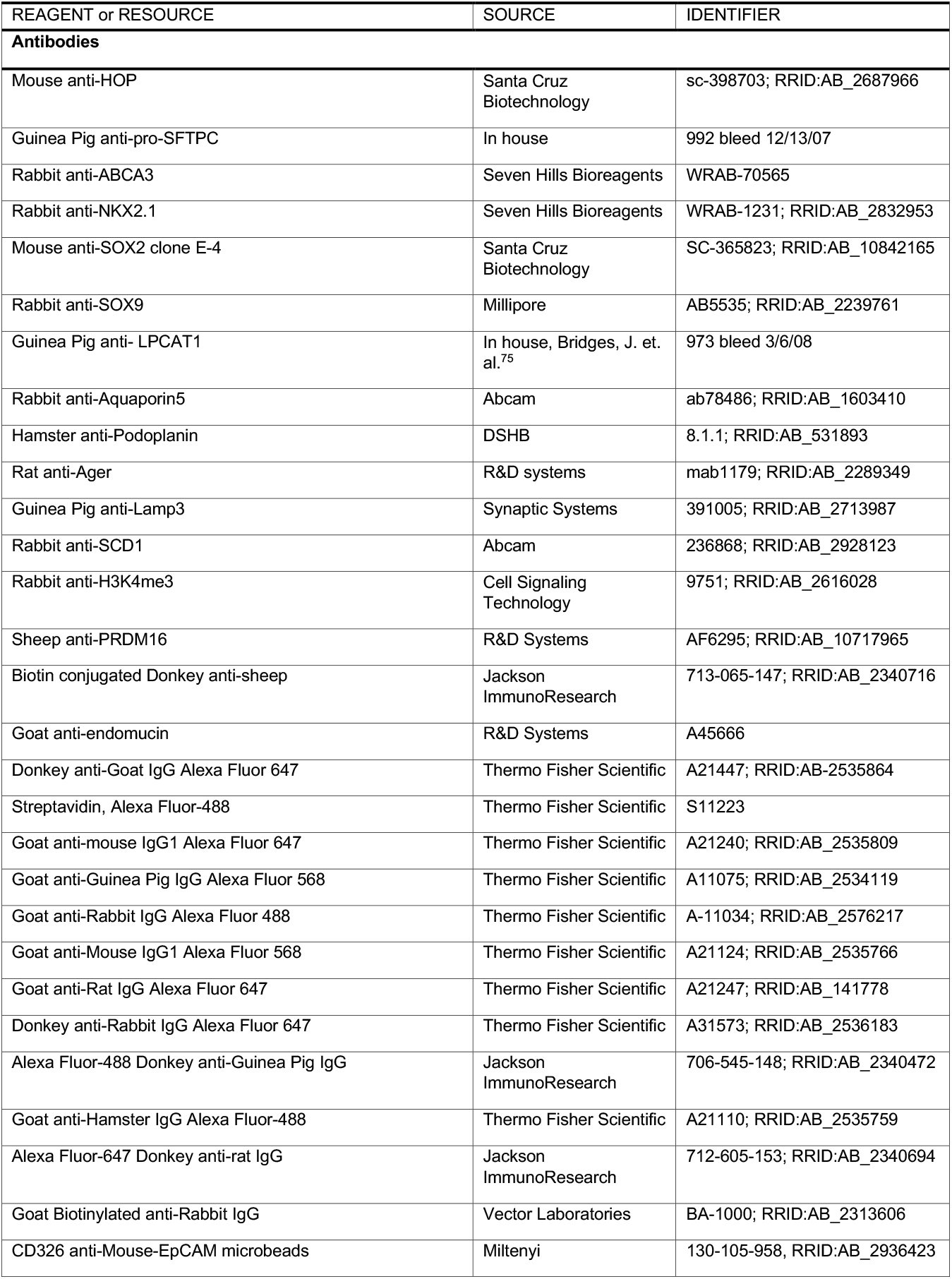
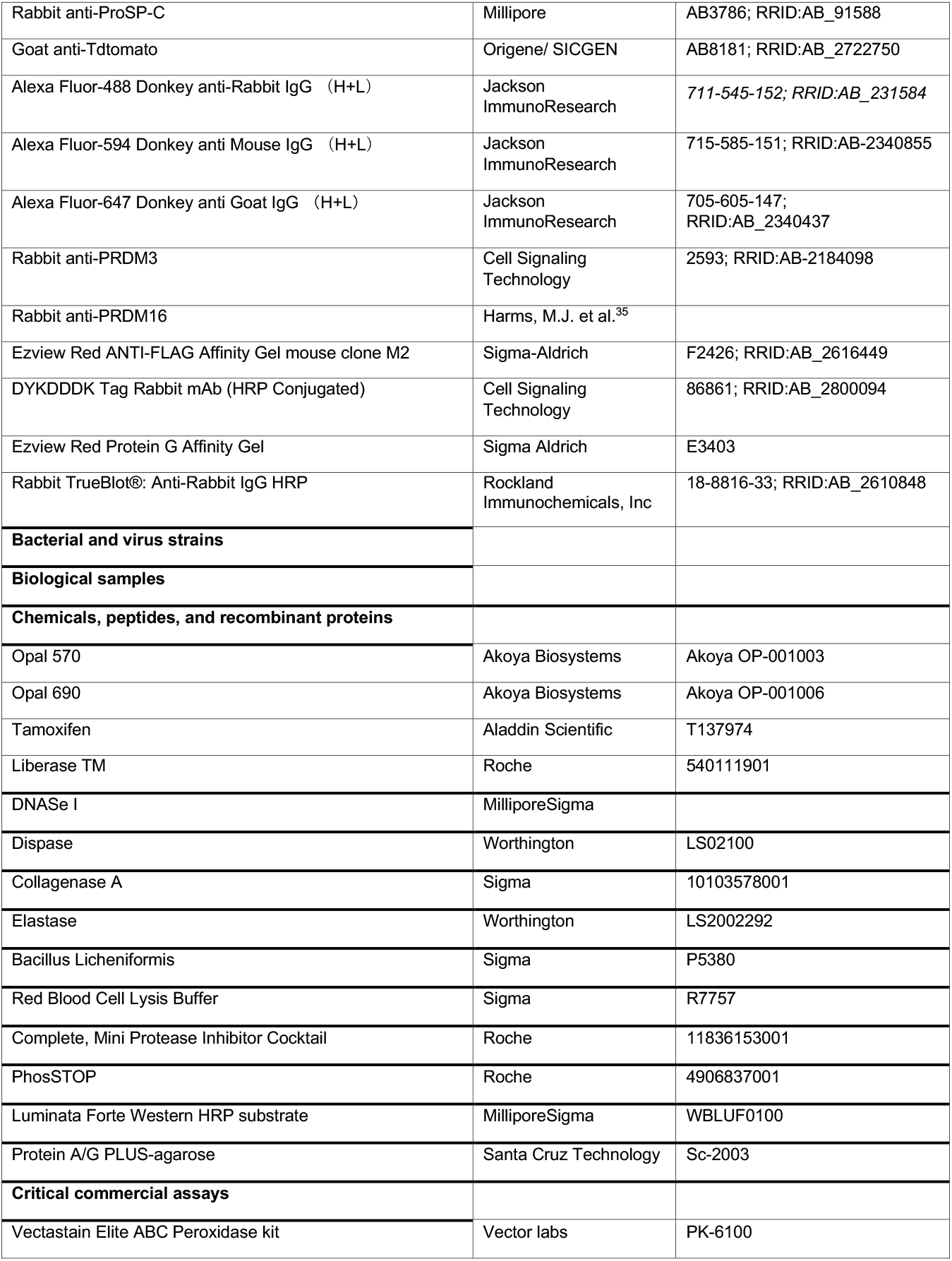
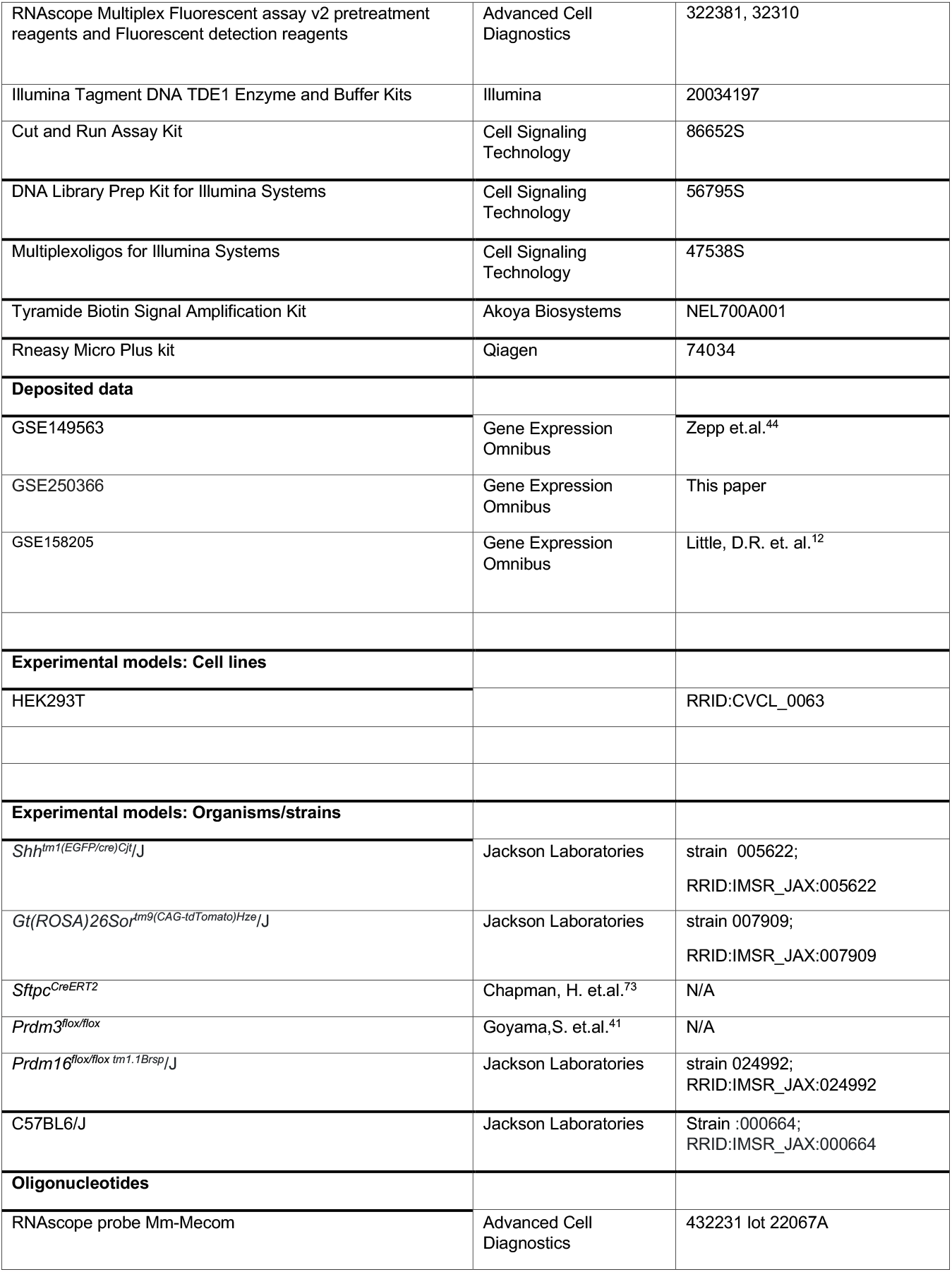
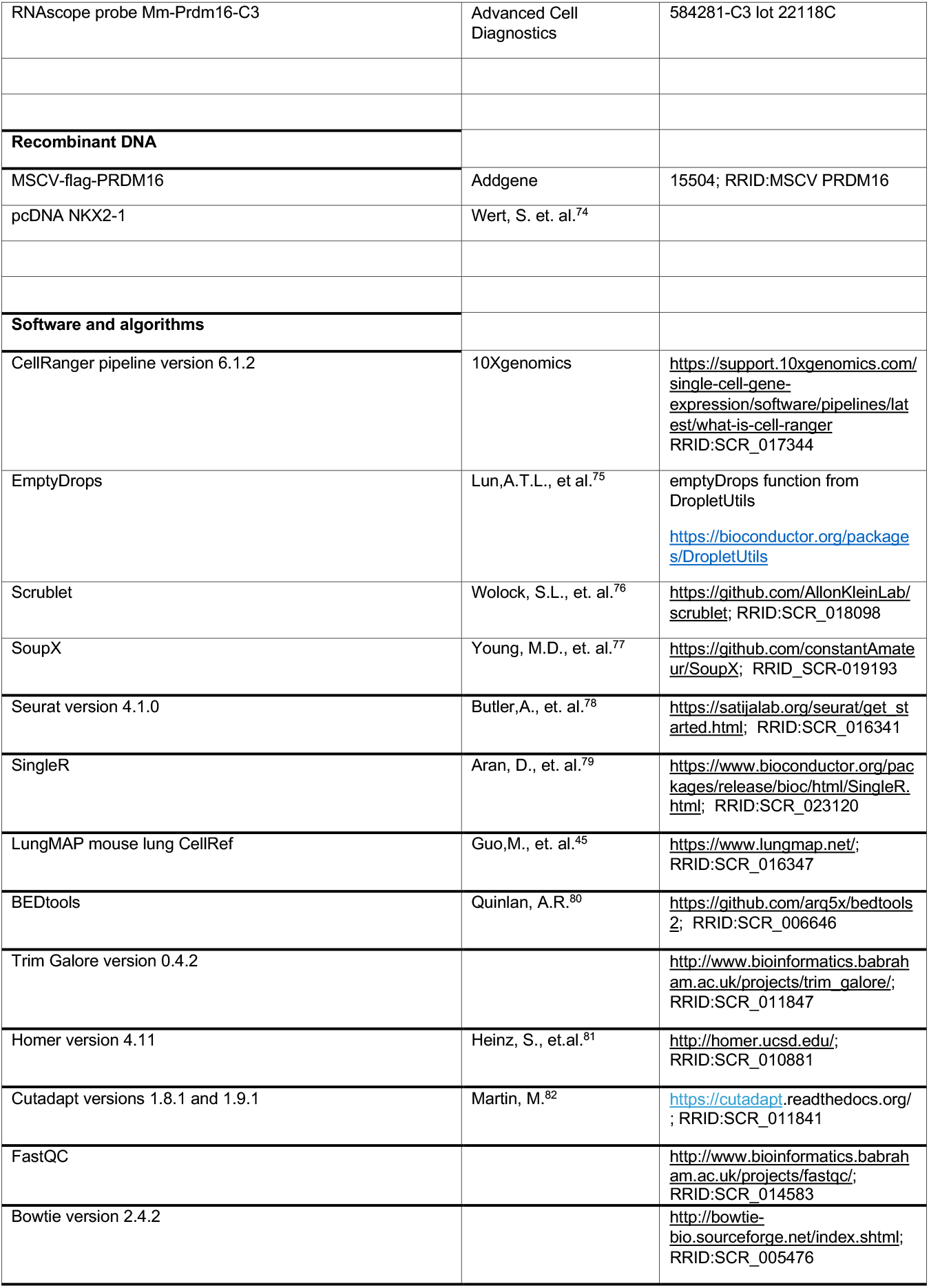
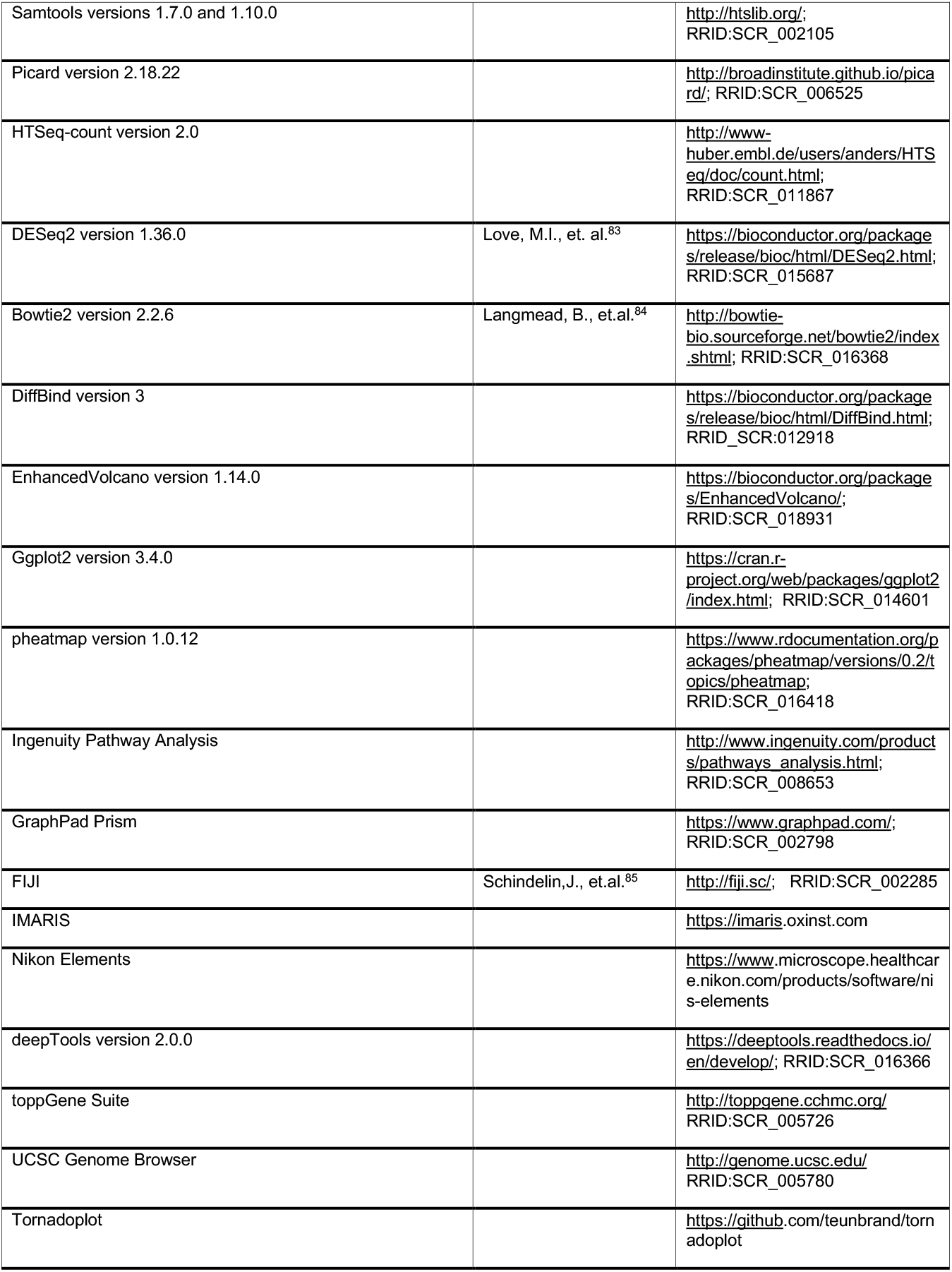
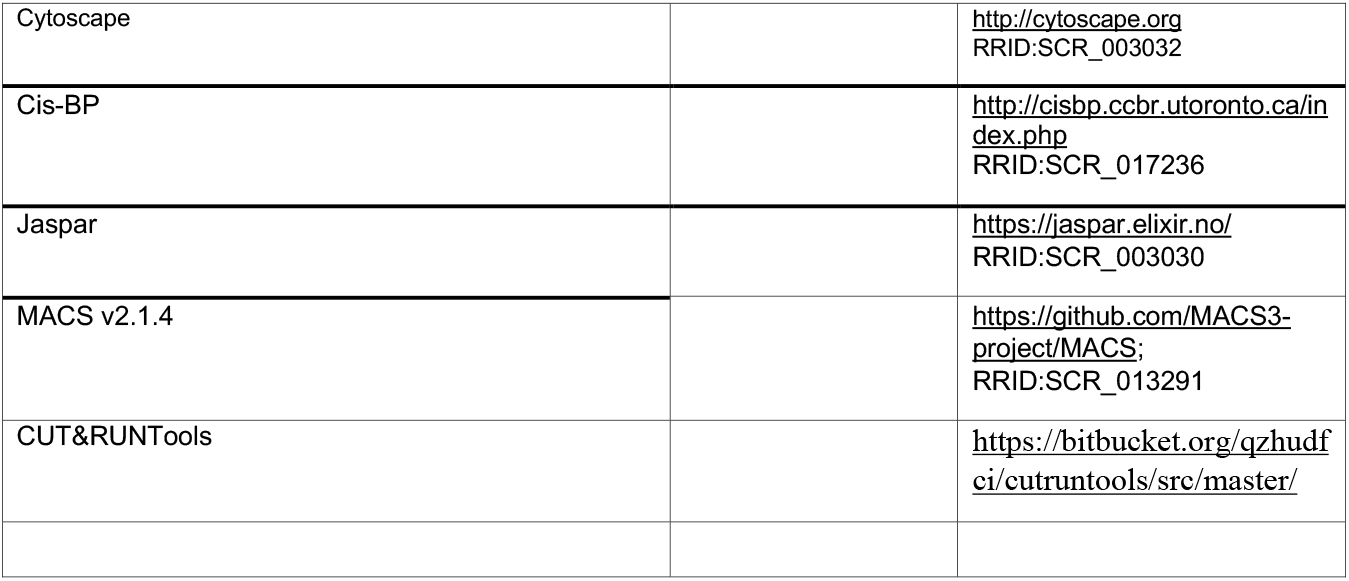

## Supplementary Information

**Figure S1:**
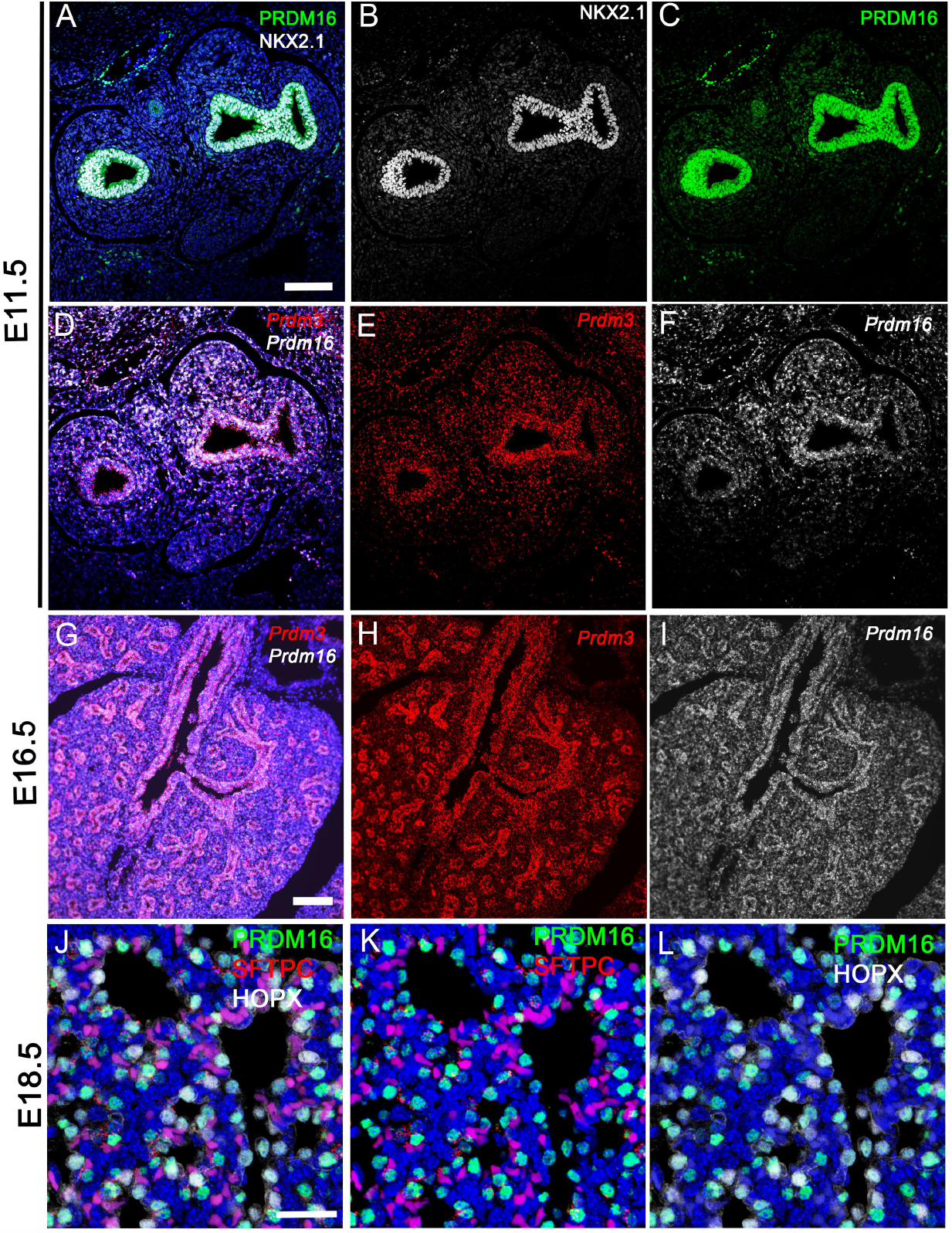
Co-ordinate expression of *Prdm3*, PRDM16, and NKX2-1 during development. A-C) Immunofluorescence of PRDM16 and NKX2-1. D-I) RNA scope analysis of Prdm3 and 16 at the indicated time points increased expression of both genes in lung epithelial cells. J-L) Immunofluorescence demonstrating PRDM16 staining in AT2 (SFTPC^+^) and AT1 (HOPX^+^) cells.

**Figure S2:**
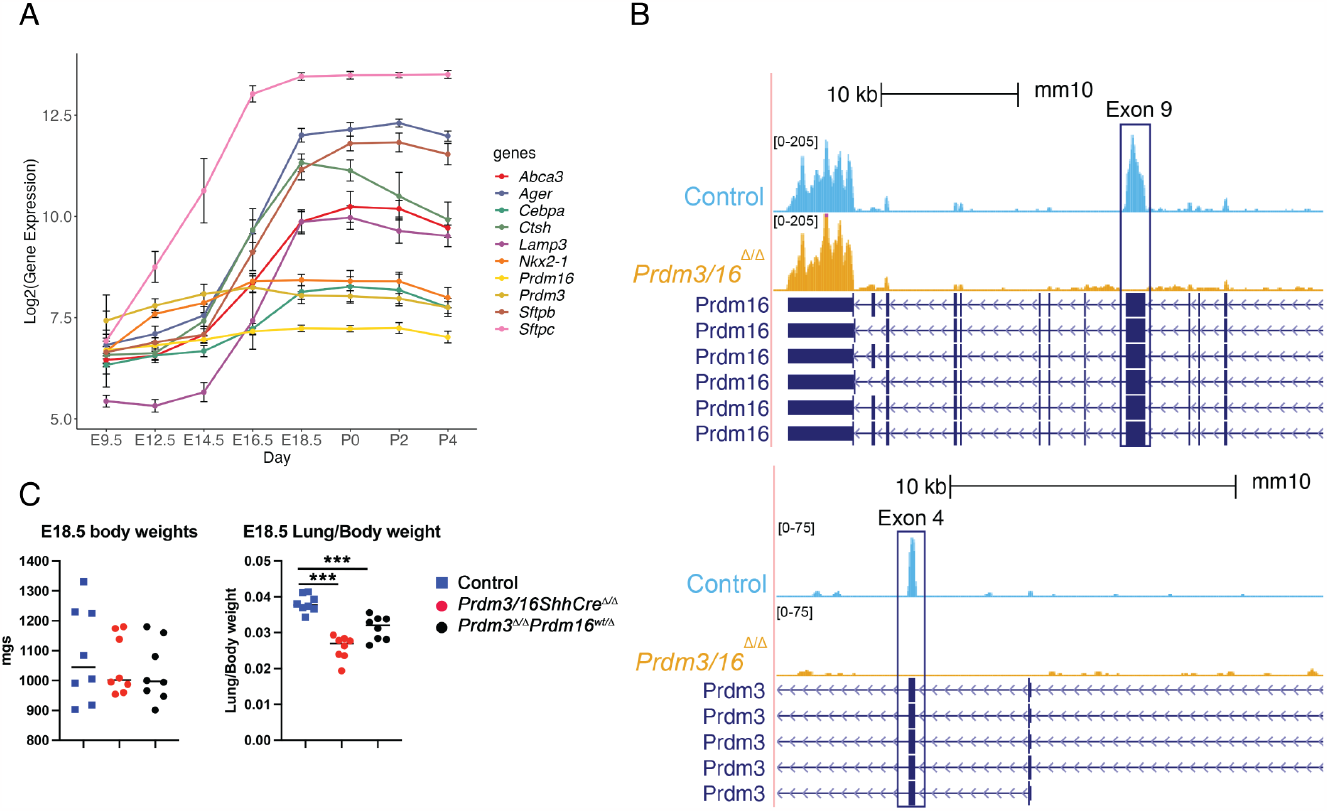
Fetal lung characteristics. A) Temporal expression of AT2 and AT1 associated RNA during lung development demonstrate relatively stable levels of *Prdm3, Prdm16*, and *Nkx2-1* and the perinatal induction of AT2 and AT1 associated RNAs, taken from Beauchemin et. al.^40^ B) Transcript analysis of the *Prdm16* and *Prdm3* loci taken from bulk RNA analysis of EpCAM+ epithelial cells demonstrates deletion of the *floxed* exons. C) Fetal whole body and lung weight data from E18.5 fetuses isolated from 5 different litters. ***p<0.0001 by 2-tailed T-test.

**Figure S3:**
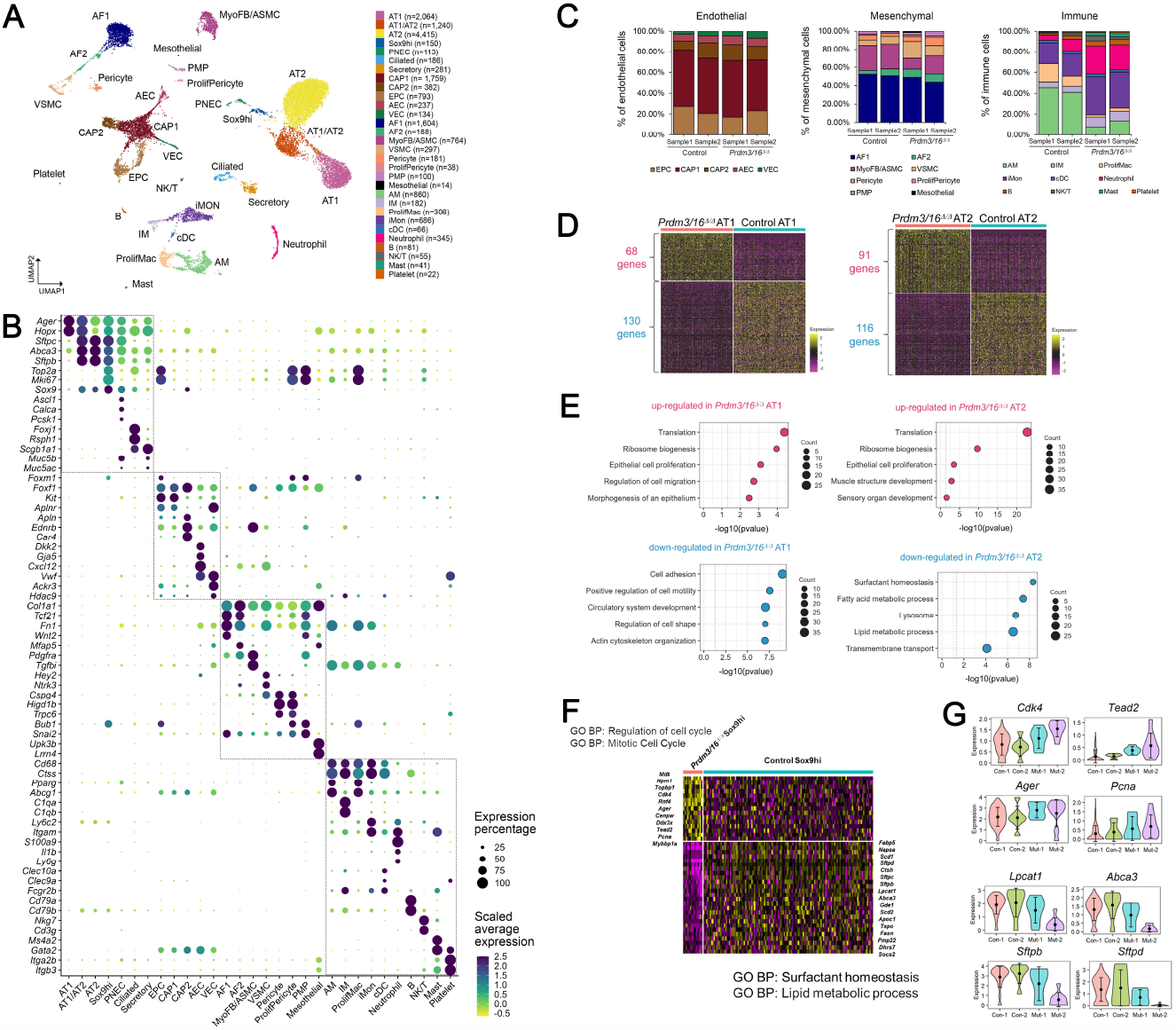
Single-cell RNA-seq (scRNA-seq) analysis of Control and *Prdm3/16*^*ShhCre*D/D^ mouse lung at E18.5. (A) UMAP plot of integrated scRNA-seq data from control (n=2 samples) and *Prdm3/16ShhCre*^Δ/Δ^ (n=2 samples) mouse lung at E18.5. Cells were colored by predicted cell types. (B) Validation of cell type prediction using expression of cell type marker genes. Dotplot visualization showed the z-score-scaled average expression level (dot color) and percentage (dot size) of each gene in each cell type. (C) Endothelial, mesenchymal, and immune cell type proportions in each of the scRNA-seq samples. (D) Differential expression analysis comparing *Prdm3/16*^*Δ/Δ*^ and control AT1 and AT2 cells using single cell RNA-seq. Representative heatmap visualization of z-score scaled expression of differentially expressed genes (DEGs) in *Prdm3/16*^*Δ/Δ*^ vs. control AT1 (left) and AT2 (right) cells from scRNA-seq of mouse lung at E18.5. For AT1, 1000 cells were randomly sampled for the visualization (500 cells from each of the *Prdm3/16*^*Δ/Δ*^ and control AT1). For AT2, 500 cells were randomly sampled for the visualization (250 cells from each of the *Prdm3/16*^*Δ/Δ*^ and Control AT2). The random sampling was repeated 10 times. All 10 heatmap visualizations showed consistent differential expression patterns. DEGs were defined based on the following criteria: p-value of Wilcoxon rank sum test <0.05, fold change >=1.5, and expression percentage >=20%. (E) Enriched Gene Ontology Biological Process annotations by the DEGs in (D). Enrichment was performed using the ToppGene suite using the DEGs as input. (F) Heat map of DEGs from the *Sox9* high epithelial population demonstrates decreased expression of AT2 cell markers and increased AT1 and proliferation markers. (G) Violin plots of selected differentially expressed genes within the Sox9 high population.

**Fig S4.**
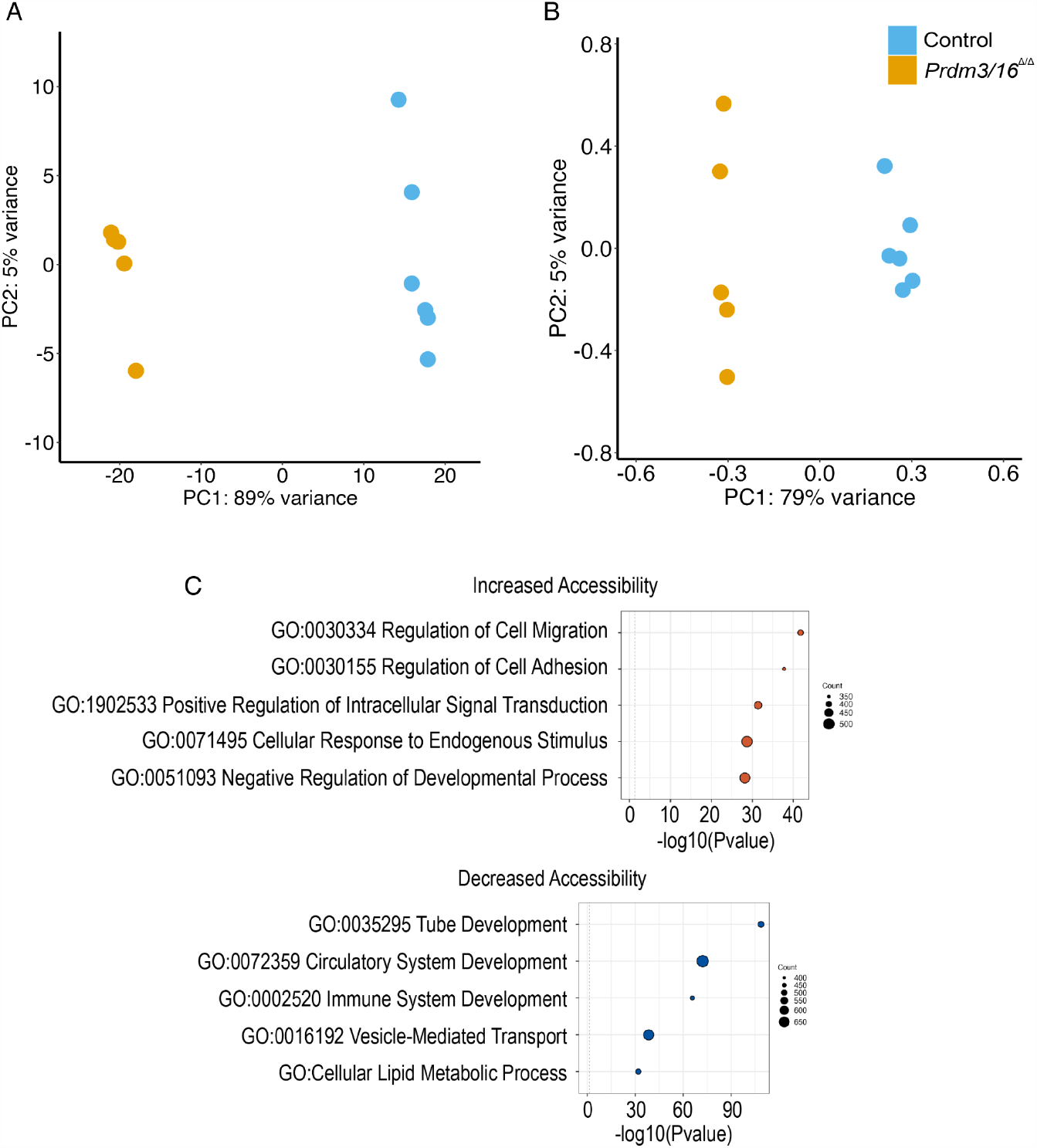
Bulk and ATAC analysis of control and *Prdm3/16*^*ShhCreΔ/Δ*^ Epcam+ sorted epithelial cells. (A) PCA of RNA sequencing of individual samples at E17.5. (B) PCA of individual ATAC sequencing. (C) Functional enrichment of genes associated with regions of increased or decreased accessibility using the tool GREAT (Genomic Regions Enrichment of Annotations Tool) is shown.

**Figure S5:**
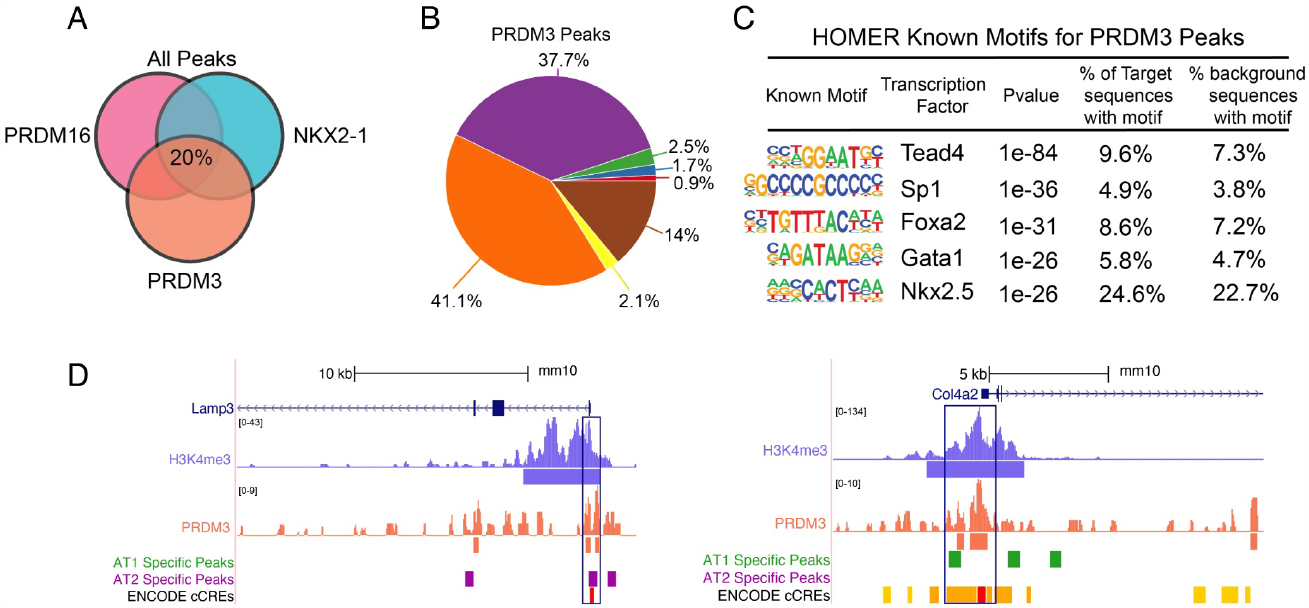
Analysis of PRDM3 CUT&RUN data from control EpCAM^+^ epithelial cells at E17.5. (A) Venn Diagram shows the overlap of PRDM3 CUT&RUN (n=1), PRDM16 CUT&RUN (4 replicates across two independent experiments), and NKX2-1 CUT&RUN (3 replicates across two independent experiments) (B) Genomic distributions of all called peaks for PRDM3 CUT & RUN are shown. (C) HOMER motif enrichment for all called peaks across the genome. (D) CUT&RUN binding is visualized with the UCSC Genome Browser for H3K4me3 and PRDM3 at the *Lamp3* and *Col4a2* loci. Sites of AT2 and AT1 cell-associated peaks from Little et al.^12^ are denoted as are the cCRE as defined in ENCODE.

**Figure S6:**
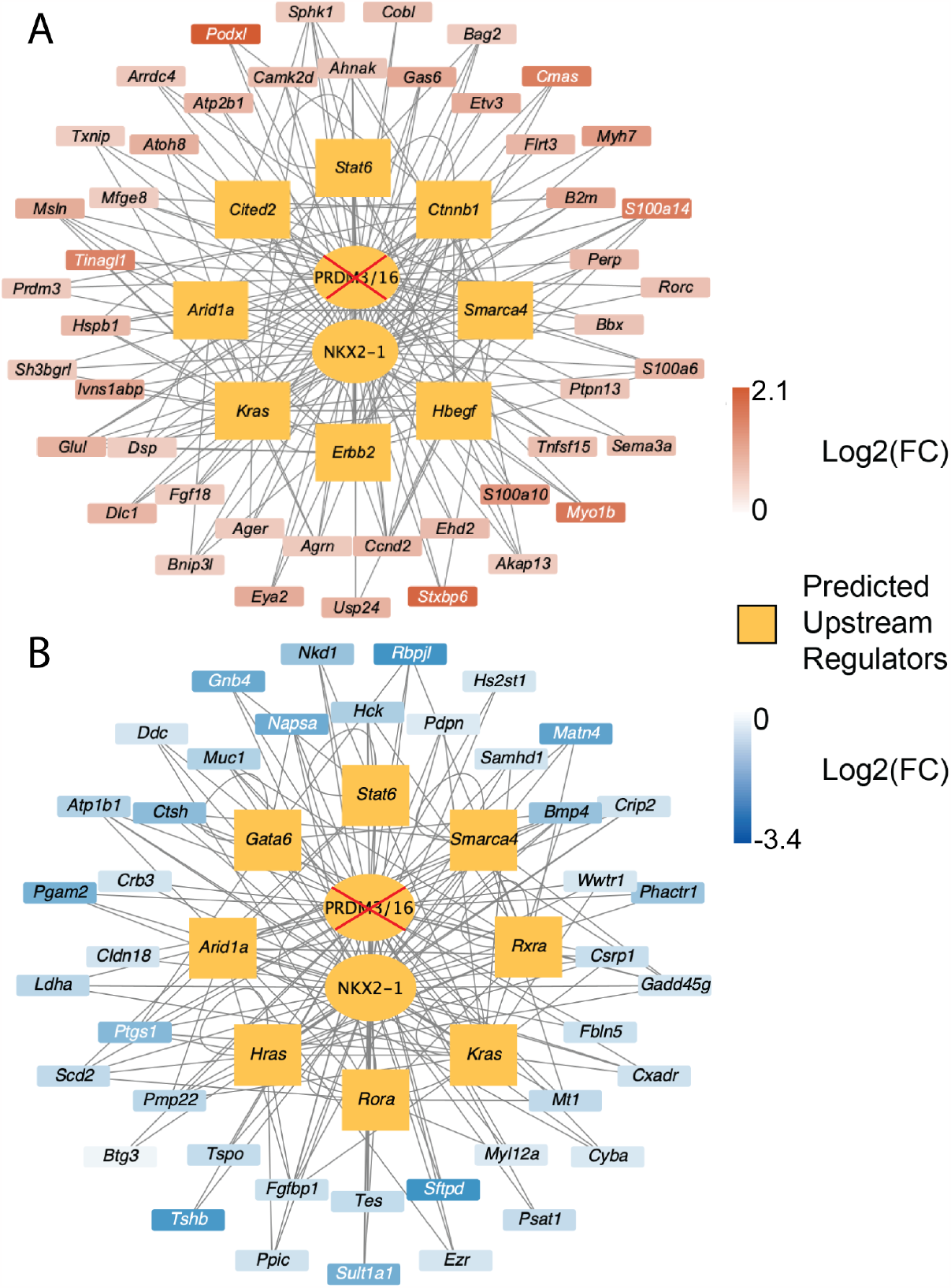
IPA analysis of differentially expressed AT1 cell associated genes. Gene regulatory networks for AT1 cells with PRDM16 and NKX2-1 at the center. Genes were selected based on the differential expression for bulk RNAseq (log2Fold change > |0.58|, p-val <0.05) and either differentially expressed in the single cell DEA or single cell gene expression (>20% expression in AT1 cell population and *Prdm3/16*^*ΔΔ*^ expression > Control expression). Those genes were used as input into IPA for a Regulatory Upstream Analysis. The resulting relationships were loaded into Cytoscape. A.) Predicted upstream regulators of genes upregulated in AT1 cells at E17.5 and E18.5. B.) Predicted upstream regulators of genes down regulated in AT1 cells at E17.5 and E18.5. Note that ARID1A, SMARC4A, KRAS, and STAT6 are predicted to modulate AT1 gene expression in both directions.

## Notes

### Competing Interest Statement

The authors have declared no competing interest.

